# Molecular Basis of EWS Interdomain Self-Association and Its Role in Condensate Formation

**DOI:** 10.1101/2025.04.16.649245

**Authors:** Erich J. Sohn, Kandarp A. Sojitra, Leticia Rodrigues, Xiaoping Xu, Bess Frost, Jeetain Mittal, David S. Libich

**Author notes:** Corresponding Authors: **David S. Libich, PhD**, Department of Biochemistry and Structural Biology, Greehey Children’s Cancer Research Institute, UT Health at San Antonio, 8403 Floyd Curl Drive, San Antonio, TX 78229-9840, **Jeetain Mittal, PhD**, Artie Mc Ferrin Department of Chemical Engineering, Texas A&M University, 200 Jack E. Brown Engineering Bldg., College Station, TX 77843-3122. These authors contributed equally (Erich J. Sohn, Kandarp A. Sojitra).

## Abstract

Ewing sarcoma, the second most common pediatric bone and soft tissue cancer, is caused by aberrant fusion of the EWS low-complexity domain (EWS^LCD^) to the DNA-binding domain of the transcription factor FLI1. The resulting fusion, EWS::FLI1, directly interacts with and engages in a dynamic interplay with EWS that drives tumorigenesis and regulates the function of both proteins. While EWS^LCD^ is known to promote self-association, the role of the RNA-binding domains (RBDs) of EWS, which include RGG repeat regions and a structured RNA-recognition motif (RRM), remains less well understood. Here, we investigate the interplay between EWS^LCD^ and RBDs using biomolecular condensation assays, microscopy, NMR spectroscopy, and molecular simulations. Our studies reveal that RBDs differentially influence EWS^LCD^ condensate formation and suggest that electrostatics and polypeptide-chain length likely contribute to this interaction. NMR spectroscopy and molecular dynamics simulations further demonstrate that EWS^LCD^ and the central RNA-binding region, comprising the RRM and RGG2 domains, engage in transient, non-specific interactions that are broadly distributed across both regions and involve diverse residue types. Specifically, tyrosine, polar residues, and proline within EWS^LCD^ preferentially interact with arginine, glycine, and proline residues in the RBD. Atomistic simulations of EWS confirm that the full-length protein exhibits a similar interaction profile with conserved chemical specificity, supporting a model in which a network of weak, distributed interdomain contacts underlies EWS self-association. Together, these findings provide molecular insight into the mechanisms of EWS condensate formation and lay the groundwork for understanding how interdomain interactions regulate EWS and EWS::FLI1 function.

**SIGNIFICANCE:** This study shows that the RNA-binding regions of the RNA-binding protein EWS subtly tune biomolecular condensation through numerous transient contacts with the low complexity domain that is common to both EWS and the oncogenic fusion EWS::FLI1. These results provide insight into condensate formation of EWS, which may be useful for understanding the oncogenic mechanisms of EWS::FLI1 and broadly, into pathogenic biomolecular condensate formation.

## INTRODUCTION

Ewing sarcoma is the prototypical example of a family of related cancers caused by chromosomal translocations involving the FET family of RNA-binding proteins, comprised of Fused in Sarcoma (<u>F</u>US), RNA binding protein EWS (<u>E</u>WS), and TATA-binding protein-associated factor 2N (<u>T</u>AF15) (Turc-Carel et al. 1983; Delattre et al. 1992; Flucke et al. 2021). The most common of these translocations fuses 264 residues from the low complexity domain (LCD) of EWS (EWS^LCD^) to the DNA-binding domain of friend leukemia integration 1 (FLI1) (Fisher 2014; Flucke et al. 2021; Maki et al. 2022). The resulting oncofusion, EWS::FLI1, is found in ∼85% of Ewing sarcoma tumors, and its ability to form biomolecular condensates is required for oncogenic transformation (Lessnick et al. 1995; Bertolotti et al. 1998; Jaishankar et al. 1999; Ng et al. 2007; Johnson et al. 2017). Biomolecular condensation of EWS and EWS::FLI1 occurs via favorable interactions within and between EWS^LCD^ domains (Spahn et al. 2003; Ng et al. 2007; Ahmed et al. 2021; Nosella et al. 2021; Johnson et al. 2024). These interactions promote EWS::FLI1 binding to gene enhancers (Johnson et al. 2017; Chong et al. 2018; Zuo et al. 2021; Vasileva et al. 2024) and recruit transcriptional co-factors (Petermann et al. 1998; Kwon et al. 2013; Boulay et al. 2017; Kim et al. 2024), promoting subsequent oncogenic transactivation.

Wild type EWS and EWS::FLI1 are co-expressed in affected individuals as only one of two EWS loci carry the aberrant fusion. Expression of the EWS::FLI1 fusion protein exerts a dominant negative repression over wild type EWS function, resulting in a phenotype that mimics EWS knockout (Jaishankar et al. 1999; Yang et al. 2000; Embree et al. 2009; Gorthi et al. 2018; Boone et al. 2021). Loss of endogenous EWS, as well as EWS::FLI1 expression, leads to poly [ADP-ribose] polymerase 1 (PARP1) accumulation at sites of DNA damage and subsequent sensitivity to DNA damaging agents (Li et al. 2007; Gorthi et al. 2018; Lee et al. 2020). Furthermore, EWS has an antagonistic effect on the activity of EWS-oncofusions, with homotypic EWS^LCD^-EWS^LCD^ interactions repressing transactivation (Chong et al. 2022), as well as heterotypic interactions between the RNA-binding domains (RBDs) of EWS and the EWS^LCD^ (Li and Lee 2000; Alex and Lee 2005). Understanding the molecular mechanisms driving EWS interdomain interactions is crucial for elucidating the interplay between EWS and EWS::FLI1.

In this work, we investigate interdomain interactions between the EWS^LCD^ and RBDs and their ability to drive EWS condensate formation using turbidity and condensate partitioning assays, microscopy, nuclear magnetic resonance (NMR) titrations, and molecular simulations. We find a strikingly heterogeneous effect of the three arginine-glycine-glycine (RGG) repeat domains on EWS^LCD^ condensate formation that mirrors their sequential and compositional heterogeneity. Extensive physiochemical characterization of condensate-forming properties between EWS^LCD^ and the central RNA-binding region, which includes RGG2, reveals electrostatics (charge interactions) and EWS^LCD^ polypeptide chain length as key drivers of the interdomain association. Using a fragment-based NMR approach we probed the atomistic details of the transient association between these two regions in the dilute phase. These experiments, along with atomistic simulations, uncover numerous non-specific contacts across both polypeptide chains that drive the interdomain self-association. In silico studies confirm that our fragment-based approach effectively recapitulates the interactions that occur in full-length EWS. Our analysis establish the structural underpinnings of full-length EWS condensate formation, and offer a basis for exploring the molecular relationship between EWS::FLI1 and wild type EWS that drives Ewing sarcoma oncogenesis.

## RESULTS

### Interdomain contacts occur throughout full-length EWS

The EWS protein largely lacks secondary structure, with the entire SYGQP-rich LCD (residues 1-280) and 67% of the RBD being intrinsically disordered (Figure S1a). The only folded domains are the RNA-recognition motif (RRM) and zinc finger (ZnF), both of which contribute to nucleic acid binding (Figure S1a) (Selig et al. 2023; Lay et al. 2024). Within the central RNA-binding region, EWS^RRM-RGG2^, *R*_1_, *R*_2_, and hetNOE measurements reporting on ps-ns time-scale dynamics are consistent with the RRM adopting a folded structure, while the RGG2 remains dynamic and disordered under these conditions (Figure S1b). Intrinsic disorder within the C-terminal RBD is due to the three RGG domains, of which RGG3 is the longest (90 residues), and RGG2 (66 residues) and RGG1 (59 residues) are somewhat similar in length (Figure 1a and S1a). RGG3 has a canonical arginine-glycine-glycine repeat structure, containing 12 RGG repeats, comprising 40% of the domain, with arginine and glycine accounting for 63% of all residues (Figure 1a). RGG1 and RGG2 are more divergent, containing only 6 and 4 RGG repeats, comprising 31% and 18% of each domain, respectively. Notably, RGG1 carries a net negative charge due to an abundance of acidic residues, particularly aspartate, while RGG2 is enriched (23%) in proline residues (Figure 1a). Sequence charge decoration, an order parameter that predicts charge patterning in intrinsically disordered regions (Sawle and Ghosh 2015), predicts RGG1 has the most charge segregation (−3.046), with negative charges clustered at the C-terminus.

**Figure 1.**
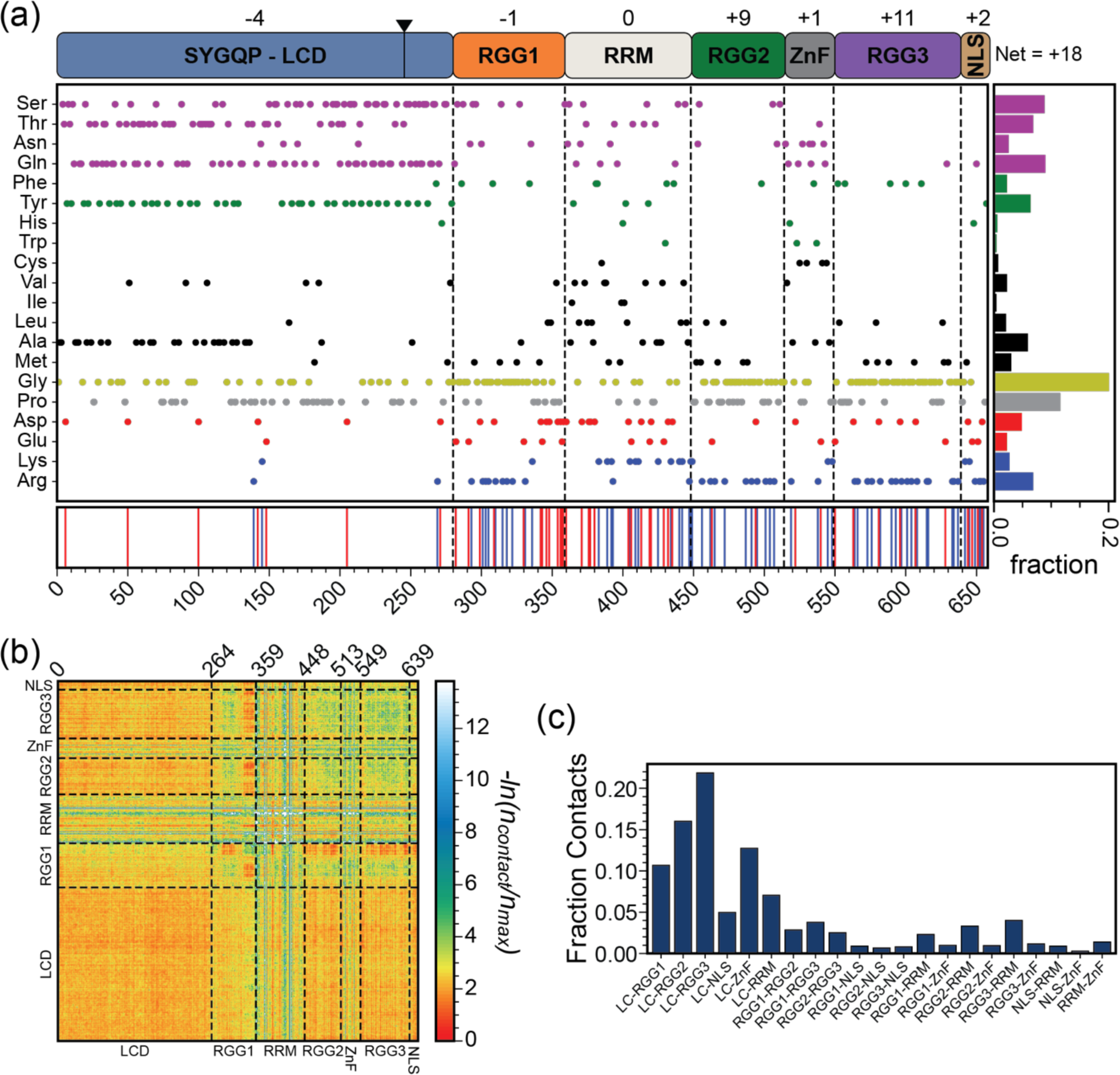
EWS^LCD^ forms interdomain contacts with RBD motifs. (a) EWS domain architecture, net charge of each domain, and amino acid distribution. The EWS::FLI1 breakpoint (264) is marked with a black triangle; red and blue lines are negatively and positively charged residues. (b) Per-residue interchain contact map and (c) fraction of total EWS interdomain contacts from co-existence CG simulations of EWS.

To investigate how the sequence features of full-length EWS governs the interdomain interactions within the condensed phase we performed coarse-grained co-existence simulations which revealed frequent contacts between the LCD and RBD (Figure 1b). While the most frequent contacts occur between the LCD and RGG domains, we detect contacts with the folded RRM and ZnF as well as between the negative charged region in RGG1 and other RGGs (Figure 1b,c). These interdomain interactions appear to enhance the condensate forming propensity of the protein, as full-length EWS has a simulated saturation concentration (C_sat_) that is ∼10 fold lower than EWS^LCD^ (Figure S2a). Based on net charge, one might expect full-length EWS to exhibit stronger electrostatic repulsion compared to EWS^LCD^ (residues 1-264), potentially reducing its phase separation (Figure 1a). However, the opposite behavior is observed, suggesting that the multivalency inherent to the longer polypeptide chain of EWS as well as charge patterning may contribute to promoting phase separation (Figure S2a). Overall, these results clearly demonstrate the propensity for the LCD to form highly promiscuous intramolecular interactions throughout the entire polypeptide chain, and especially with the three RGG domains.

### RBDs of EWS impact EWS^LCD^ condensate formation

To investigate the role of interdomain contacts in EWS condensate formation, we created two-component systems combining EWS^LCD^ with EWS^RGG1-RRM^, EWS^RRM-RGG2^ and EWS^RGG3^ (Figure S2b,c). In a single-component system, both EWS^LCD^ and EWS::FLI1 form condensates through diverse interaction modes, stimulated by dissolved ions (salting out), consistent with the effect of other LCDs (Murthy et al. 2019; Martin et al. 2020; Bock et al. 2021; Krainer et al. 2021; Johnson et al. 2024).

In the absence of NaCl, EWS^LCD^ and EWS^LCD^ + EWS^RGG1-RRM^ solutions have virtually no turbidity and lack visible condensates, consistent with previous reports for the EWS^LCD^ (Nosella et al. 2021) under these conditions (Figure 2a,b). In the presence of 150 mM NaCl, addition of EWS^RGG1-RRM^ diminished solution turbidity compared to EWS^LCD^ control, indicating reduced condensate formation, despite maintaining similar morphology as EWS^LCD^-only condensates (Figure 2a,b). EWS^RRM-RGG2^ and EWS^RGG3^ both substantially increased EWS^LCD^ condensate formation in the absence of NaCl, although EWS^RGG3^ had a greater effect (Figure 2a). Upon addition of 150 mM NaCl, EWS^LCD^ + EWS^RRM-RGG2^ and EWS^LCD^ + EWS^RGG3^ condensate formation was diminished, with the EWS^LCD^ + EWS^RGG3^ samples having a smaller change (Figure 2a). In the presence of NaCl, EWS^LCD^ + EWS^RRM-RGG2^ condensate morphology resembles that of EWS^LCD^-only samples, however, in the absence of NaCl they appear less round (Figure 2b). Condensates formed by EWS^LCD^ + EWS^RGG3^ were a mixture of small and spherical, which remained suspended, and irregularly shaped, which appeared to settle quickly, spreading out heterogeneously across the slide, regardless of whether NaCl was present (Figure 2b).

**Figure 2.**
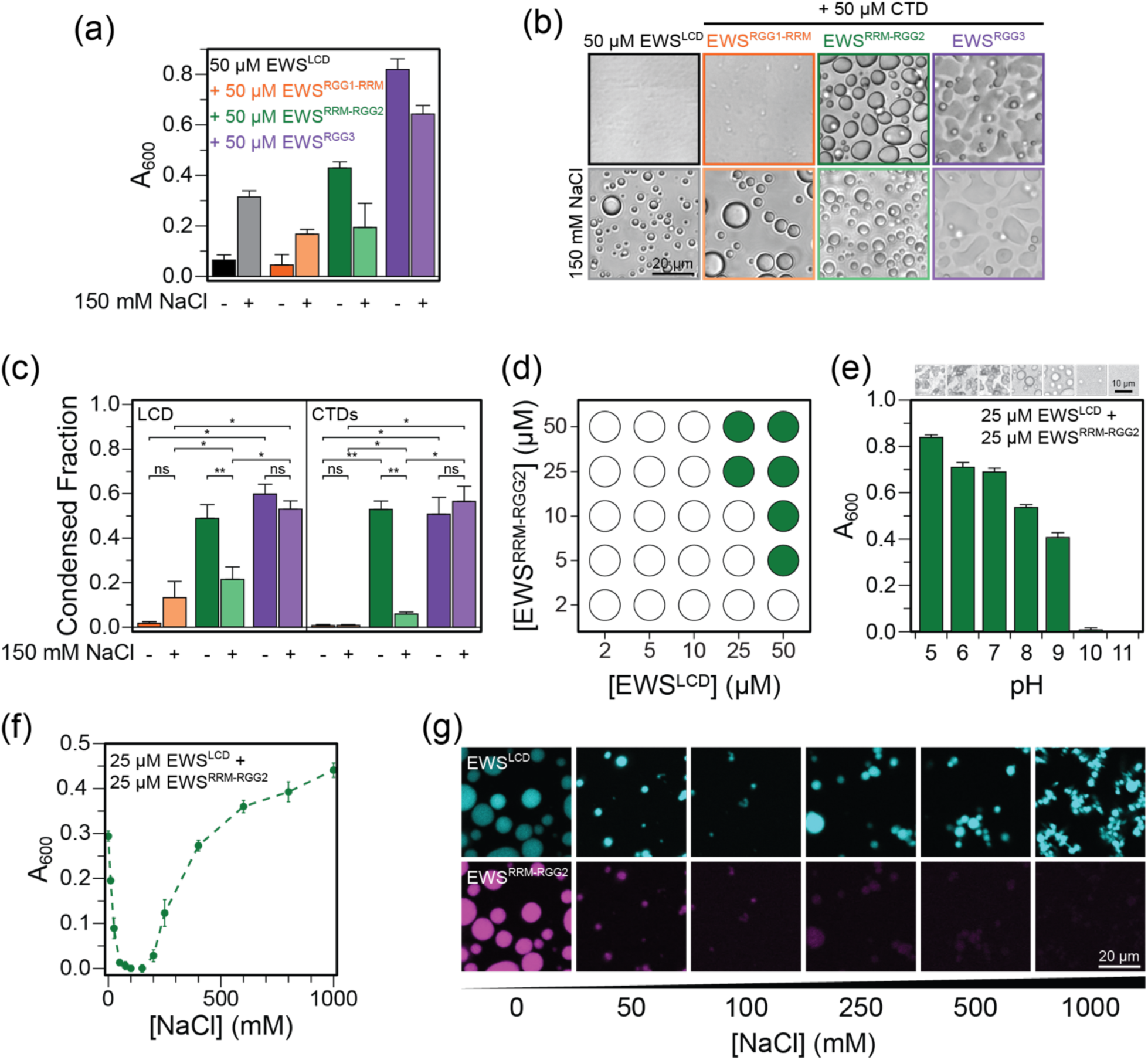
EWS RBD motifs impact condensate formation propensity of EWS^LCD^. (a) Turbidity measurements, (b) bright field microscopy, scale bar = 20 µm, and (c) condensate partitioning of 50 μM EWS^LCD^ only (black) and + 50 μM of EWS^RGG1-RRM^ (orange), EWS^RRM-RGG2^ (green), or EWS^RGG3^ (purple). Lighter shade is with 150 mM NaCl. (d) Phase diagram for EWS^LCD^ + EWS^RRM-RGG2^ (filled circles = visible condensates). Turbidity measurements of EWS^LCD^ + EWS^RRM-RGG2^ across different (e) pH, scale bar = 10 µm, and (f) NaCl concentrations. (g) Fluorescence microscopy of select [NaCl] from (f), scale bar = 20 µm. Error bars = SD.

Partitioning assays were used to gain insight into the distinct effects of each RBD on EWS^LCD^ condensate formation. In EWS^LCD^ + EWS^RGG1-RRM^ samples, both domains remained completely soluble in the absence of NaCl, consistent with turbidity measurements (Figure 2c). Addition of NaCl had no effect on the solubility of EWS^RGG1-RRM^, and only ∼13% w/w of EWS^LCD^ was driven into condensates (Figure 2c). Conversely, ∼53% w/w EWS^RRM-RGG2^ and ∼51% w/w EWS^RGG3^ co-partitioned into condensates with approximately equal amounts of EWS^LCD^ (60% and 49% w/w, respectively) in the absence of NaCl (Figure 2c). Addition of 150 mM NaCl significantly reduced EWS^RRM-RGG2^ (∼6% w/w) and EWS^LCD^ (21% w/w) condensate co-partitioning, while insignificantly changing the amounts of EWS^RGG3^ (∼56% w/w) and EWS^LCD^ (∼53% w/w) that co-partition. The unique impact of each RBD constructs on EWS^LCD^ condensates suggests a complex interplay of interdomain interactions that drive EWS self-association that are regulated through distinct biochemical properties.

To further investigate the intriguing response of the EWS^LCD^ + EWS^RRM-RGG2^ system to NaCl, we created a phase diagram varying concentrations of EWS^LCD^ and EWS^RRM-RGG2^ (Figures 2d and S3a). In a single component system absent of NaCl, the C_sat_ of EWS^LCD^ is > 150 μM (Johnson et al. 2024). Addition of equal molar ratios of EWS^RRM-RGG2^ reduced the EWS^LCD^ C_sat_ to < 25 μM. When EWS^RRM-RGG2^ is present in molar excess, condensates form in solution at < 10 μM EWS^LCD^ (Figures 2d and S3a). Similarly, 50 μM EWS^LCD^ was induced to form condensates by < 5 μM EWS^RRM-RGG2^ (Figures 2d and S3a). To investigate the role of multivalency, we reduced the EWS^LCD^ length by fragmenting it into three overlapping regions: EWS^1-120^, EWS^91-199^, and EWS^171-264^ (Figure S2b,c) (Johnson et al. 2022). Surprisingly, none of the three fragments formed condensates with EWS^RRM-RGG2^ when mixed at equimolar ratios up to 150 μM (Figure S3b), suggesting that multivalency and length are critical factors driving the association.

EWS^LCD^ + EWS^RRM-RGG2^ condensate formation was more complex than the initial turbidity and condensate partitioning assays revealed. While equimolar ratios (25 μM) of EWS^LCD^ and EWS^RRM-RGG2^ readily form condensates from pH 5 to 9, formation is inhibited at pH 10 and 11 (Figure 2e). In the absence of NaCl, robust turbidity is measured for the bi-component EWS^LCD^ + EWS^RRM-RGG2^ system. Upon addition of up to 50 mM NaCl, turbidity is reduced (Figures 2f and S3c). At moderate NaCl concentrations (50 - 200 mM), condensate formation is almost completely inhibited, yet as NaCl increases further (> 200 mM), turbidity again increases, indicating the reappearance of condensates (Figures 2f and S3c). Fluorescent microscopy images at selected NaCl concentrations reveal that, EWS^LCD^ and EWS^RRM-RGG2^ colocalize into condensates in the absence of NaCl and that low levels of NaCl in solution greatly reduce the number and size of the condensates (Figure 2g). However, at higher NaCl concentrations, the condensates that reappear almost entirely exclude EWS^RRM-RGG2^ and form smaller fusion-defective condensates (Figure 2g), consistent with previous findings of single-component EWS^LCD^ condensates at high NaCl concentrations (Johnson et al. 2024). These two distinct modes of condensate formation reveal the complexity of EWS self-association and the ability of EWS to drive molecular organization across a wide range of physiochemical conditions. The ability of soluble ions to prevent EWS^LCD^ and EWS^RRM-RGG2^ from co-localizing into condensates suggests a potential role of electrostatic interactions in driving co-condensation of these two domains.

### Atomistic details of association between ^15^N-EWS^LCD^ and EWS^RRM-RGG2^

We used NMR and molecular simulations to investigate the association and dynamics of the dilute phase interaction between EWS^LCD^ and EWS^RRM-RGG2^ as a surrogate for full-length EWS. To abrogate pH changes and dilution effects we used a “reverse” titration protocol of the ^15^N-EWS^LCD^ fragments with EWS^RRM-RGG2^ (Dcosta et al. 2024), and observed numerous small chemical shift perturbations (CSPs) dispersed across the sequence of each fragment (Figures 3 and S4a). Minor line broadening was also observed for residues across the three fragments, yet no obvious clustering indicative of a single binding site is evident (Figure S4b). EWS^1-120^ had the greatest proportion of both shifted and broadened peaks, suggesting that this region contains a higher density of interaction sites with EWS^RRM-RGG2^ (Figures 3a and S4b). Both CSPs and line broadening were small in magnitude and non-sequential, suggesting that EWS^LCD^ non-specifically interacts with EWS^RRM-RGG2^ with minor contributions from residues throughout the polypeptide chain (Figure 3 and S4b). Consistent with these NMR observations, dilute phase atomistic simulations of EWS^LCD^ fragments with EWS^RRM-RGG2^ (Figure S5a-c) also revealed non-specific contacts distributed across the lengths of the EWS^LCD^ fragments (Figure 3). Among the three EWS^LCD^ fragments, EWS^1-120^ appeared to form more contacts with EWS^RRM-RGG2^ compared to the other fragments (Figure 3a). Notably, while residues within the 40-60 region of EWS^1-120^ exhibited higher contact frequencies, the interactions remained broadly distributed throughout the LCD, highlighting the overall non-specific nature of these interactions (Figure 3a).

**Figure 3.**
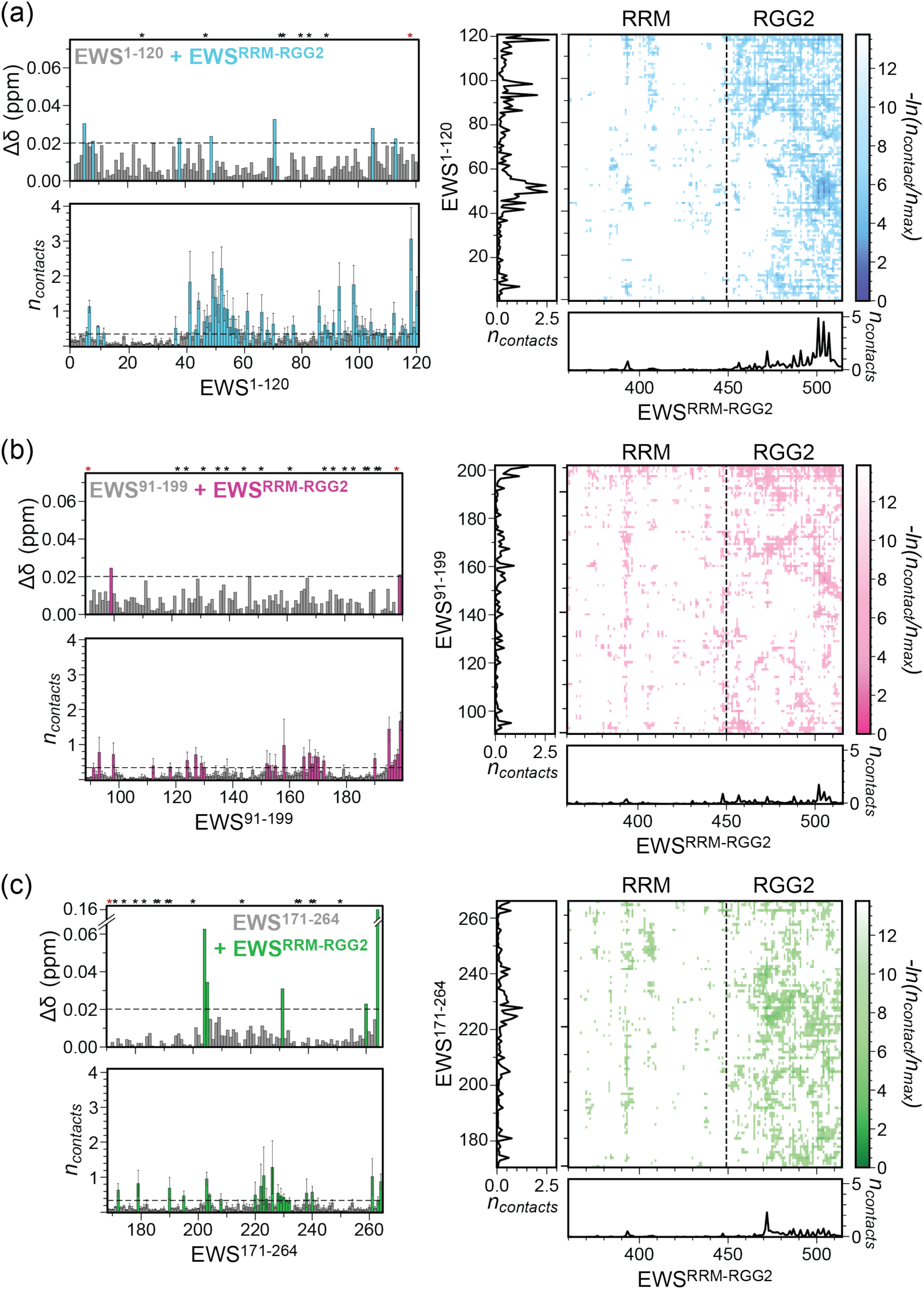
EWS^LCD^ interacts with EWS^RRM-RGG2^ through contacts distributed across their chains. Chemical shift perturbations (CSPs) (top left panels), average per-residue contacts (bottom left panels), and normalized average pairwise van der Waals contacts (right panels) for (a) ^15^N-EWS^1-120^ (cyan), (b) ^15^N-EWS^91-199^ (pink), and (c) ^15^N-EWS^171-264^ (green) interacting with EWS^RRM-RGG2^. CSPs are the maximum observed after addition of 3:1 molar ratio of EWS^RRM-RGG2^ and color-filled if CSPs > 1 SD (dashed black line). Asterisks indicate overlapped/ambiguous assignments (red) or prolines (black). Average per-residue and pairwise van der Waals contacts were calculated from atomistic simulations of EWS^RRM-RGG2^ with each EWS^LCD^ fragment and are colored filled if contact values are > the mean for all three fragments (dashed black line). The average per-residue contacts formed between EWS^LCD^ fragments and EWS^RRM-RGG2^ are shown along the left and bottom margins of each contact map.

### Atomistic details of association between ^15^N-EWS^RRM-RGG2^ and EWS^LCD^

We next performed the inverse titration of ^15^N-EWS^RRM-RGG2^ with EWS^LCD^ fragments. Since only 39% of non-proline residues of RGG2 are assignable due to sequence degeneracy and spectral overlap, we stratified the unassignable peaks as glycine or arginine/other based on their ^13^C chemical shifts (Selig et al. 2023) (Figure 4). The unassigned peaks nearly all come from RGG2, as 98% of all non-proline residues in the RRM are assigned, and thus this approach enabled tracking of RG, RGG, and PGG motifs even without sequence specific assignments. Titration of ^15^N-EWS^RRM-RGG2^ with each of the three EWS^LCD^ fragments resulted in small CSPs and minimal line broadening for residues distributed in both the RRM and RGG2 regions (Figures 4 and S6). As with the ^15^N-EWS^LCD^ titrations, CSPs were small in magnitude and dispersed across the EWS^RRM-RGG2^ polypeptide chain (Figure 4), supporting the conclusion that the interdomain association is largely non-specific with minor contributions from numerous residue pairs. The increased chain flexibility provided by the sequence disorder found in RGG2 may permit the increased contact propensity with the intrinsically disordered EWS^LCD^ fragments (Figure S1). Similarly, dilute phase atomistic simulations of EWS^RRM-RGG2^ with EWS^LCD^ fragments reveals mostly non-specific contacts distributed across the length of RRM and RGG2 save for EWS^RRM-RGG2^ residues 390-395, which interacts with all three fragments and is part of a surface exposed loop within the RRM (Figure S7a,b). Moreover, the RGG2 domain displayed a higher number of interactions with EWS^LCD^ fragments compared to the RRM (Figures 3 and S7a,b).

**Figure 4.**
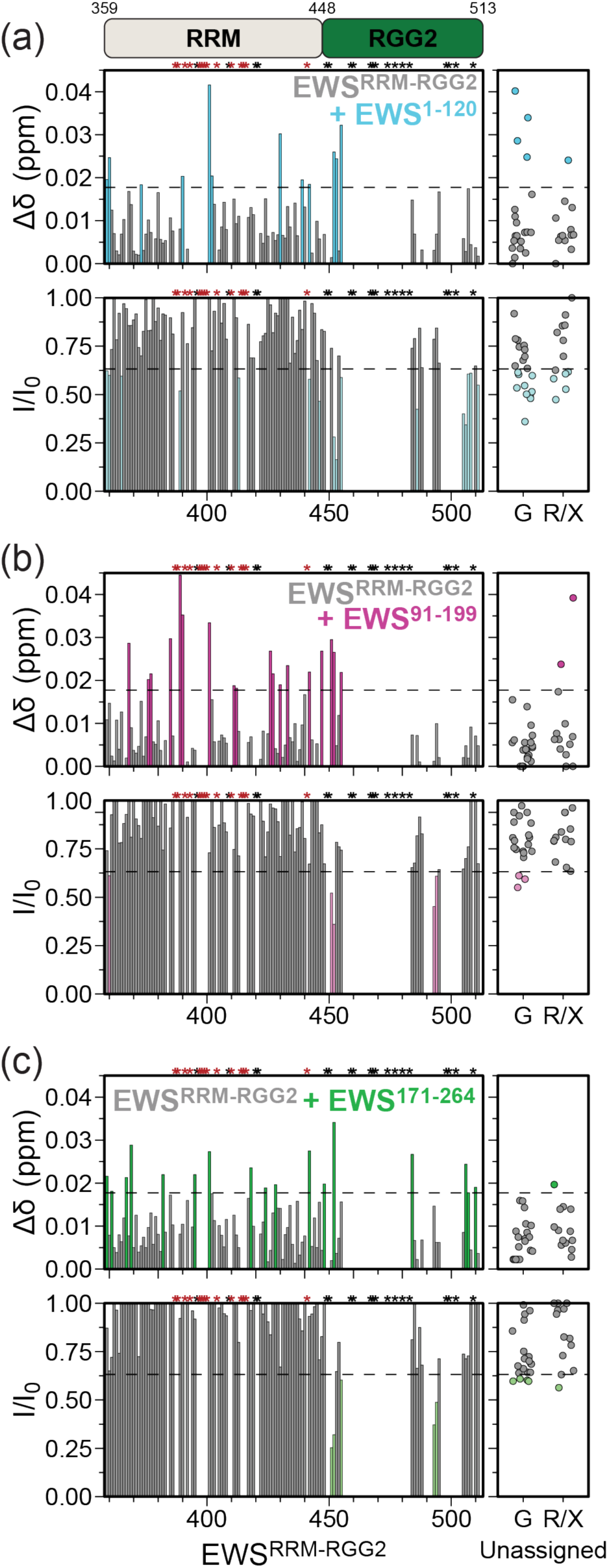
RRM and RGG2 form interdomain contacts with EWS^LCD^ fragments. CSPs (top panels) and line broadening (bottom panels) of ^15^N-EWS^RRM-RGG2^ titrated with (3:1 molar ratio) (a) EWS^1-120^, (b) EWS^91-199^, or (c) EWS^171-264^. CSPs (line broadening) > (<) 1 SD (dashed line) are plotted in cyan (EWS^1-120^), pink (EWS^91-199^), or green (EWS^171-264^). Asterisks mark unassigned/ambiguous/broadened residues (red) and proline (black). Unassigned glycine (G) or unclassified (R/X) resonances are plotted (right panels).

Several EWS^RRM-RGG2^ residues consistently shifted and/or broadened with the different EWS^LCD^ fragments, suggesting that residue types and the local environment mediate interactions. Residues N452 and R455 were shifted in three and two titrations, respectively, and two unassigned non-glycine residues were broadened in two of the three titrations (Figure S7c). Additionally, M451, N452, D493, and R494 peaks were all broadened by all three EWS^LCD^ fragments, representing two pairs of neighboring residues that made consistent contact regardless of which region of EWS^LCD^ was present (Figure S7c). CSPs and line broadening did not localize to a specific region of EWS^RRM-RGG2^ RRM domain (Figure S8a-d) and were dispersed between both the positively-charged nucleic acid binding face and the opposite negatively-charged face (Figure S8e). Analysis of simulation data reveal K391-R392 and R446-K447 consistently formed interactions with the three EWS^LCD^ fragments (Figure S7a,b). Within RGG2, the sequence FPPRGPRGSR showed a higher level of interaction with all three EWS^LCD^ fragments. These interactions remained transient, as reflected by fluctuations in contact probabilities (Figure S7b).

### Numerous residue types contribute to the interaction between EWS^LCD^ and EWS^RRM-RGG2^

We stratified the observed CSPs and line broadening from EWS^LCD^ titrations to serine, tyrosine, glycine, glutamine, proline, alanine, and other (these residues comprise 93% of the sequence), and those from EWS^RRM-RGG2^ by residue type (Figure S9a,b). Shifts occurred in nearly 40% of the “other” residues within EWS^LCD^, indicating that this interaction is driven by a wide variety of non-specific contacts (Figure S9a). Peak shifts were detected in each type of residue within EWS^RRM-RGG2^, with 36% of all residues shifted by at least one of the three EWS^LCD^ fragments, >60% of positively charged residues, and >40% of polar residues shifted (Figure S9b). Analysis of residue contributions from dilute phase simulations revealed that various residue positions in both EWS^LCD^ fragments and EWS^RRM-RGG2^ form transient contacts. In EWS^LCD^ fragments, tyrosine, polar residues, proline, and aspartate contributed to interactions with EWS^RRM-RGG2^ (Figure S9c). In EWS^RRM-RGG2^, arginine, phenylalanine, polar residues, proline, and methionine appear to interact with the EWS^LCD^ fragments, with arginine, glycine and proline being prominent contributors (Figure S9c).

Together, these results suggest that the LCD or RBD lack a single specific binding region that mediates their association. Instead, the interdomain association is stabilized by an accumulation of relatively weak, but highly numerous contacts between a wide variety of residue pairs that are dispersed across the entire EWS^LCD^ and both RRM and RGG2. In the full EWS^LCD^, these interactions become numerous enough to be potent drivers of condensate formation, which are likely to be even more readily formed in full-length EWS, where additional contacts are made between the LCD and the other RBDs.

### Interdomain association between EWS^LCD^ and EWS^RRM-RGG2^ in full-length EWS

To understand how interdomain interactions manifest in the context of full-length EWS, we employed dilute-phase atomistic simulations (Figure S5d). Over the course of 16 µs, the LCD of full length EWS interacted with parts of the RBD, preferring disordered segments over the folded domains, and the folded RBD elements predominantly interacted with their flanking regions (Figure 5a). Comparing the simulations of full-length and EWS^LCD^ fragments revealed that the overall pattern of interactions between EWS^LCD^ and EWS^RRM-RGG2^ are consistent (Figure 5b). An interesting deviation emerged in EWS between residues 150–200, (breakpoint between EWS^99-199^ and EWS^171-264^) which showed reduced interactions with EWS^RRM-RGG2^ in the full-length protein relative to the EWS^LCD^ fragments (Figure 5b). This segment preferentially interacted with other RBDs, such as RGG1 and RGG3, suggesting that in the full-length system, structural constraints restrict conformational flexibility, while the presence of additional RBDs provides alternative interaction partners, thereby limiting its ability to engage with RRM-RGG2 (Figure 5b). In contrast, in isolated fragment simulations, this region was unconstrained and able to explore a broader conformational ensemble, allowing stronger interactions with EWS^RRM-RGG2^ (Figure 5b). Indeed, inter-residue distances between the EWS^LCD^ fragments in their isolated state versus the full-length protein revealed that EWS^LCD^ fragments were more collapsed in the full-length system (Figure S10a). Further, residue-type analysis revealed consistent patterns of interactions between the full-length and fragment simulations identifying tyrosine, polar residues, proline, and aspartate (EWS^LCD^), and arginine, proline, and glycine (EWS^RRM-RGG2^) as the key residues contributing to the interaction network (Figures 5c, and S10b). Representative snapshots from full-length EWS simulations further support the transient and non-specific nature of these interactions (Figure 5d). Together, these results provide confidence that our biophysical studies of simplified systems capture the transient, non-specific interactions that are relevant to self-association and phase separation in the context of full-length EWS.

**Figure 5.**
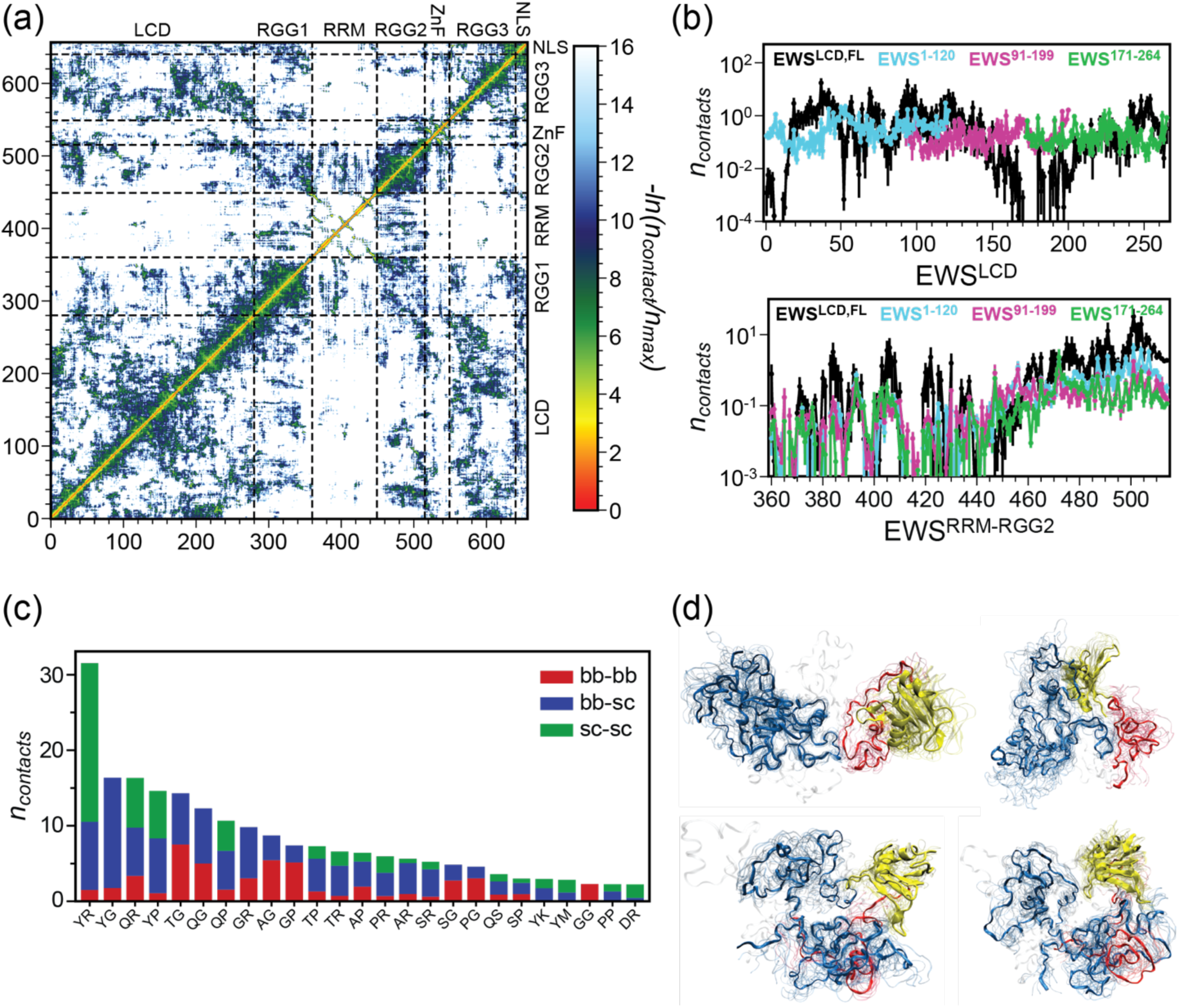
RRM and RGG2 form interdomain contacts with the LCD in full-length EWS. (a) Normalized average pairwise van der Waals contacts from single chain atomistic simulations of full-length EWS. (b) Average per-residue contacts formed by with EWS^RRM-RGG2^ (top) or by EWS^RRM-RGG2^ with EWS LCD constructs (bottom). EWS^LCD,FL^ (extracted from atomistic simulations of single chain full-length EWS, black), EWS^1-120^ (cyan), EWS^91-199^ (pink), or EWS^171-264^ (green). (c) Average pairwise interdomain residue type contacts between EWS^LCD^ and EWS^RRM-RGG2^ extracted from single chain atomistic simulations of full-length EWS; backbone-backbone (red), backbone-sidechain (blue), and sidechain-sidechain (green), first residue is from EWS^LCD^, second residue is from EWS^RRM-RGG2^. (d) Examples of EWS^LCD^-EWS^RRM-RGG2^ ensemble images (EWS^LCD^, blue; RRM, yellow; RGG2, red) from atomistic simulations of full-length EWS.

## DISCUSSION

Many of the functions of EWS and EWS::FLI1 are thought to be mediated through condensate formation, that is primarily driven by inter- and intra-molecular interactions between multiple residue types within the EWS^LCD^ (Kato et al. 2012; Kwon et al. 2013; Chong et al. 2018; Nosella et al. 2021; Johnson et al. 2024; Sundara Rajan et al. 2024). However, studies have also demonstrated interdomain interactions between the LCD and RBD of FUS (and other RNA-binding proteins) are important for condensate formation (Martin et al. 2021; Murthy et al. 2021; Wake et al. 2025). Here we uncover similar, transient, non-specific interactions between the EWS LCD and RBDs, and discover that charge and sequence patterning within the RGG motifs differentially influence EWS phase separation. Our findings are consistent with the growing body of work pointing to both the patterning of interaction sites and the quantity of contacts, rather than specific location as key drivers of biomolecular condensation (Lin et al. 2017; Martin et al. 2020; Galagedera et al. 2023; Rekhi et al. 2023; Rekhi et al. 2024).

Recent studies on miscibility of intrinsically disordered proteins (IDPs) highlighted that client partitioning is governed by the balance between homotypic and heterotypic interactions between the client and scaffold proteins (Rana et al. 2024; Welles et al. 2024). Moreover, studies have highlighted that charge segregation enhances compaction and condensate formation (Das and Pappu 2013; Sawle and Ghosh 2015; Schuster et al. 2020). Therefore, our observations of reduced co-partitioning of EWS^RGG1-RRM^ compared to EWS^RRM-RGG2^ and EWS^RGG3^ with EWS^LCD^ may arise from sequence differences between the RGG motifs. In addition to having weaker heterotypic interactions with EWS^LCD^, the charge patterning in RGG1 may increase its homotypic strength (keeping it below the threshold for phase separation), allowing it to compete with heterotypic interactions, while heterotypic interactions of EWS^RRM-RGG2^ and EWS^RGG3^ with EWS^LCD^ appear sufficient to support co-partitioning under similar conditions. Co-partitioning can also be aided by homotypic interactions within the client domains, provided that heterotypic interactions are sufficiently strong (Welles et al. 2024). Notably, upon addition of salt, partitioning of both EWS^RRM-RGG2^ and EWS^RGG3^ into condensates is reduced, likely due to weakened electrostatic interactions that disrupt both homotypic and heterotypic contacts. However, condensate partitioning assays and imaging tell a more complex story, wherein salt addition only significantly affects the condensates with EWS^RRM-RGG2^. While EWS^LCD^ condensates remain spherical over time, addition of EWS^RRM-RGG2^ or EWS^RGG3^ causes non-spherical droplet morphologies, an effect more pronounced with EWS^RGG3^. This greater surface-wetting effect may be due to the greater positive charge carried by EWS^RGG3^. However, in our previous work, the surface wetting effect was not detected with relatively inflexible poly-L-lysine peptides, suggesting that chain flexibility likely also plays an important role in how the proteins in a condensate interact with surfaces.

Kamagata et al. recently reported a size-dependent condensate exclusion effect in FUS droplets, where the partitioning of folded proteins is restricted while disordered domains are preferentially recruited (Kamagata et al. 2022). The retention of the folded RRM on EWS^RRM-RGG2^ may thus limit its incorporation into condensates relative to the fully disordered EWS^RGG3^. Additionally, the relative lengths and net-charge differences between the EWS RGG domains affect their condensate partitioning and influence condensate dynamics. For example, the distinct two-phase regime governed by ionic strength in the EWS^LCD^ + EWS^RRM-RGG2^ condensates highlights the sophisticated regulatory factors governing condensate formation and the highly complex interplay between the different domains of EWS. We observe a notable enrichment of contacts involving tyrosine, polar residues, proline and aspartate in EWS^LCD^ with arginine, glycine, and proline in EWS^RRM-RGG2^, suggesting a chemically selective interaction profile involving π–π interactions, hydrogen bonding, and electrostatics. Simulations identify a region within RGG2 (FPPRGPRGSR) in which arginines occur at regular intervals, with proline positioned nearby. Given that proline can engage in CH-π (Zondlo 2013) and CH-O (Daniecki et al. 2022) interactions and impart local conformational rigidity (Perez et al. 2014; Mateos et al. 2020; Hazra et al. 2023), this arrangement may support favorable local contacts and interaction dynamics. Our atomistic simulations of full-length EWS reveal that while interaction patterns change in the context of the full-length protein, with the LCD gaining more opportunities to interact with multiple RBDs, the overall chemical interaction specificity remains consistent with that observed in the fragment simulations.

## CONCLUSION

Our study shows EWS RBD-derived domains interact non-specifically with its LCD, that these interactions lack well-defined or structured binding sites and instead engage in weak, transient interactions involving multiple different types of residues. As in vitro condensates are defined by a complex network of multiple biomolecular partners, we expect that inclusion of nucleic acids, particularly RNA, and other relevant partners such as EWS::FLI1 will alter EWS dynamics in vivo. Indeed, the presence of the FLI1^DBD^, has a pronounced effect on EWS^LCD^ condensates, (Selig et al. 2022), effects which are likely to be altered in the presence of the EWS RBD. These findings provide the basis for understanding how EWS interdomain self-association occurs and influences condensate formation, and emphasizes the need for further investigation into the biological consequences of disrupting these interactions through mutagenesis, post-translational modification, and modulation with other biomolecules, particularly nucleic acids.

## MATERIALS AND METHODS

### Recombinant protein cloning and expression

Synthetic DNA genes were codon optimized for expression in *E. coli*, synthesized (GenScript, NJ), and cloned into modified pET expression vectors with an N-terminal His_8_-tag followed by a tobacco etch virus (TEV) protease cleavage site and verified by sequencing. For the EWS^RGG1-RRM^ and EWS^RGG3^, a maltose-binding protein (MBP) tag was included between a His_10_-tag and TEV cleavage site. Transformed BL21 Star^TM^ (DE3) cells were cultured Luria Broth (LB) at 37°C until OD_600_ ∼ 0.6-0.8, and protein expression was induced by 0.5 mM IPTG, grown for 3-5 hrs, harvested (30 min at 4,000 x *g*, 4°C), and the cell pellets were stored at −80°C. For ^15^N incorporation, an O/N 5 mL LB culture was pelleted (10 min at 1000 x *g*, 4°C), resuspended in 5 mL M9 minimal media supplemented with 1 g/L ^15^NH_4_Cl and 0.02% (w/v) yeast extract, expressed as before, using M9 in place of LB. All cultures were supplemented with 100 μg/mL ampicillin.

### Recombinant protein purification

Cells expressing EWS^LCD^, EWS^1-120^, EWS^91-199^, and EWS^171-264^ were purified as described previously (Johnson et al. 2022). Briefly, cell pellets were resuspended in 20 mM CAPS, pH 11, 8 M urea, 500 mM NaCl, 10 mM imidazole, lysed via sonication, and clarified (30 min at 45,000 x *g*). Cleared supernatant was loaded onto an IMAC column, washed with 1 M NaCl, and eluted with 300 mM imidazole. Pooled fractions were dialyzed against 50 mM Tris, pH 8.8, 1% 1’,6’-hexanediol, 2 mM DTT, TEV was added after 4 hrs and dialysis continued for 16 hrs at room temperature. Dialyzed protein was diluted two-fold into 20 mM CAPS, pH 11, 500 mM NaCl, 10 mM imidazole, clarified by centrifugation, and passed over an IMAC column. The flow-through was collected, concentrated to < 2 mL, and loaded onto a HiLoad 16/600 Superdex 75 pg size exclusion column (Cytiva, MA) equilibrated in 20 mM CAPS, pH 11. Fractions containing the purified protein were pooled and concentrated via a centrifuge concentrator to > 300 μM, aliquoted, and stored at −80 °C until use.

Cells expressing EWS^RRM-RGG2^ and MBP-tagged EWS^RGG1-RRM^ were purified as described for EWS^RRM-RGG2^ in our previous work (Selig et al. 2023). Briefly, cell pellets were lysed in 50 mM Tris, pH 8.5, 1 M NaCl, 15 mM imidazole, 1 mM TCEP, and clarified as described above. Cleared supernatant was loaded onto an IMAC column, washed with 30 mM imidazole, and eluted with 300 mM imidazole. Pooled fractions were dialyzed against 50 mM Tris, pH 8.0, 100 mM NaCl, 2 mM DTT, 0.5 mM EDTA. TEV was added after 4 hrs, and dialysis continued for 16 hrs at room temperature. Dialyzed protein was clarified by centrifugation, 1 M NaCl and 15 mM imidazole (final concentration) were added, and was passed over an IMAC column. The flow-through was collected, concentrated to < 2 mL and loaded onto a HiLoad 16/600 Superdex 75 pg size exclusion column (Cytiva, MA) equilibrated in 20 mM Tris, pH 7.2, 2 mM TCEP. Fractions containing the purified protein were pooled and concentrated via a centrifuge concentrator to > 300 μM, aliquoted, and stored at −80 °C until use.

Cell pellets containing MBP-tagged EWS^RGG3^ were lysed in 50 mM Tris, pH 8, 1 M NaCl, 15 mM imidazole, 1 mM TCEP, and clarified in the same way as EWS^RRM-RGG2^. Cleared supernatant was loaded onto an IMAC column, washed with 40 mM imidazole, and eluted with 300 mM imidazole. Pooled fractions were dialyzed against 50 mM Tris, pH 8.0, 1 mM TCEP for 6 hrs, diluted two-fold in the dialysis buffer, transferred to a 50 mL conical tube, TEV was added, and the sample was incubated for 16 hrs at room temperature. The pH was lowered to 7.5 by diluting with 20 mM Tris, pH 7, 1 mM TCEP, then 1 M NaCl and 15 mM imidazole (final concentration) were added. The solution was clarified by centrifugation, and passed over an IMAC column. The flow-through was collected, buffer exchanged into 20 mM Tris, pH 7.2, 2 mM TCEP via a spin concentrator (MilliporeSigma, MA), concentrated to > 300 μM, aliquoted, and stored at −80 °C until use. Purity of all purified proteins was determined by SDS-PAGE.

### Turbidity assays

25 μL samples were prepared in triplicate in 384-well flat bottom black plates (Greiner Bio-One, NC). Protein stocks were diluted so that the addition of the protein stock was only 2 μL. Samples were prepared to 20 mM Tris, pH 7.5 except the pH sensitivity assay where 20 mM of sodium acetate (pH 5), MES (pH 6), HEPES (pH 7), Tris (pH 8 and 9), or CAPS (pH 10 and 11) were substituted. Stock NaCl was added to the required concentration and samples were mixed thoroughly prior to adding protein stocks. EWS^LCD^ or EWS^LCD^ fragments were always added last, as none of the EWS RBDs initiate phase separation on their own, under the conditions tested. The plate was sealed using clear sealing tape (Thermo Scientific, MA), and turbidity was measured at 600 nm using a BioTek Cytation Gen 5 (Agilent, CA) or a GloMax Discover (Promega, WI) plate reader. For the NaCl sensitivity assay, turbidity was measured at 2 min intervals for 6 hours at 25°C. Images were acquired on a BioTek Cytation Gen 5 (Agilent, CA) using a 60 x objective immediately following turbidity measurement.

### Bright-field and Fluorescence Microscopy

EWS^LCD^ condensates with different EWS RBD constructs were prepared as described for the turbidity assays. For fluorescence microscopy, EWS^LCD^ and EWS^RRM-RGG2^ were labelled with DyLight 488 and 650, respectively via sortase transpeptidase, as previously described (Mao et al. 2004; Theile et al. 2013; Johnson et al. 2024), and mixed to ∼ 1 % (v/v) with their unlabeled counterparts. Samples were imaged using either an Olympus FV300 (Olympus, Japan) or a Zeiss LSM 980 (Zeiss, Germany) inverted confocal microscope and acquired simultaneously in bright field and fluorescent modes using a 20 or 40 x objective lens. Images were re-colored, and contrast was adjusted globally using FIJI (Schindelin et al. 2012).

### Condensate partitioning

Samples were prepared as described for the turbidity assays except the total sample volume was 100 μL to ensure visibility of the pelleted fraction. Freshly prepared samples were incubated at room temperature for 30 minutes and centrifuged at 14,000 x *g* for 20 mins at 20°C. The supernatant was removed, and the pellet was resuspended in 7 M urea. The supernatant (“dilute”), pellet (“condensed”), and a sample prior to centrifugation (“total”) were analyzed by SDS-PAGE. Band intensity (*I*) of EWS^LCD^ and EWS RBD constructs were measured by densitometry using FIJI (Schindelin et al. 2012). The “condensed fraction” was calculated using:

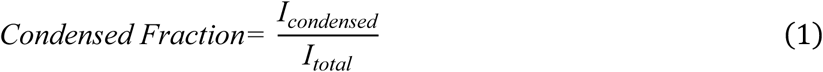

### Nuclear magnetic resonance spectroscopy

NMR experiments were collected at 298 K on a Bruker Avance NEO spectrometer (Bruker, MA) operating at a proton Larmor frequency of 700.05 MHz. Data were processed using NMRPipe (Delaglio et al. 1995) and analyzed with CCPNMR Analysis version 3.2 (Skinner et al. 2016). The ^1^H,^15^N-HSQC was recorded using 64* x 1024* complex points in the indirect (^15^N) and direct (^1^H) dimensions, corresponding to acquisition times of 33.3 and 106.5 ms, respectively, for EWS^RRM-RGG2^, and 41.0 and 112.6 ms, respectively, for the EWS^LCD^ fragments.

For the titration of EWS^LCD^ fragments with ^15^N-labelled EWS^RRM-RGG2^, 1.1 mL of 100 μΜ ^15^N-EWS^RRM-RGG2^ in 20 mM potassium phosphate buffer, pH 6, 50 mM KCl was prepared, divided into two equal aliquots. To the first aliquot 300 μM of an EWS^LCD^ fragment (final concentration) was added (saturated sample), and to the second an equivalent amount of the EWS^LCD^ fragment buffer (20 mM CAPS, pH 11) (initial sample). ^1^H,^15^N-HSQC spectra were recorded on both the “initial” and “saturated” samples. Intermediate titration points corresponding to 33.3, 50, 100, and 200 μM EWS^LCD^ fragments were obtained by mixing the “initial” and “saturated” samples at appropriate ratios to achieve a final sample of 550 μL at the desired concentration. Similarly, titration of EWS^RRM-RGG2^ with ^15^N-EWS^LCD^ fragments was as described exception the sample buffer was 20 mM MES, pH 5.5 and the “initial” sample was prepared using EWS^RRM-RGG2^ buffer of 20 mM Tris, pH 7.2, 2 mM TCEP. Spectra were processed in Topspin 4.1.1 with a sine bell window function, and zero filled to twice the number of acquired points for analysis. Chemical shift perturbations (CSP) were calculated by weighting the ^1^H and ^15^N chemical shifts with respect to their gyromagnetic ratio using:

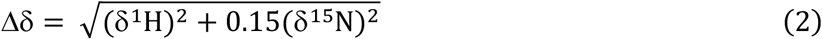

CSPs were considered significant when they were higher than the standard deviation of Δδ for all residues across the three fragments.

^15^N *R*_1_ and *R*_2_ relaxation rates were calculated from *T*_1_ and *T*_1_*_ρ_* experiments, recorded on a 200 µM sample of EWS^RRM-RGG2^ in 20 mM potassium phosphate pH 6, 50 mM KCl using 100* x 1024* complex data points in the indirect (^15^N) and direct (^1^H) dimensions corresponding to acquisition times of 43.4 and 112.6 ms, respectively. The ^15^N *T*_1_ experiment consisted of 8 interleaved spectra with the following relaxation delays: 40, 80, 120, 200, 280, 400, 600, and 800 ms. The *T*_1_*_ρ_* experiment was recorded using a *B*_1_ field of 1400 Hz and 8 interleaved spectra with the following relaxation delays: 1, 21, 31, 41, 61, 81, 121 and 161 ms. ^15^N *R*_2_ rates were calculated using the following equation:

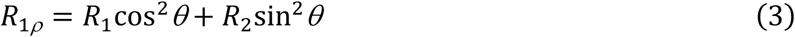

with *θ* = arctan(*ω*_1_/Ω), where *ω*_1_ is the *B*_1_ field strength (here 1400 Hz) and Ω is the offset from the spinlock carrier frequency. ^1^H-^15^N heteronuclear NOE experiments were recorded on the same sample and consisted of two interleaved experiments, with and without proton saturation, using a recycle delay of 4 seconds. Spectra were acquired with 128* x 1024* complex data points in the indirect (^15^N) and direct (^1^H) dimensions corresponding to acquisition times of 43.4 and 112.6 ms, respectively.

### Statistics

Statistical significance of differences in condensate partitioning between each EWS RBD construct added to EWS^LCD^ condensates were analysed via Student’s *t*-tests using Microsoft Excel version 16.

### Coarse-grained (CG) simulations

We simulated full-length EWS and each fragment (EWS^1-120^, EWS^91-199^, EWS^171-264^, EWS^RRM-RGG2^) using the HPS-Urry (Regy et al. 2021) CG model. Single-chain simulations were performed with LAMMPS (Thompson et al. 2022) in 30 × 30 × 30 nm cubic boxes for 2 µs. For CG co-existence simulations, we used HOOMD-blue 2.9.3 (Anderson et al. 2020) following previously established protocol (Dignon et al. 2018; Mammen Regy et al. 2021). We simulated 40 chains of full-length EWS in a 17.5 × 17.5 × 122.5 nm slab, matching a similar total protein concentration as used for EWS^LCD^ in our previous work (Johnson et al. 2024). Simulations were run with a 10 fs time step at 300 K. To compare the saturation concentrations of full-length EWS and EWS^LCD^ (Johnson et al. 2024), we performed simulations at 330 K, where full-length EWS showed coexisting phases, allowing comparison with EWS^LCD^ saturation concentrations. Temperature was controlled using a Langevin thermostat with friction coefficient γ = m_i_/t_damp_, where m_i_ is residue mass and t_damp_ was set to 1000 ps. Each simulation ran for 5 µs, with the first 1 μs excluded as equilibration time. Error bars were calculated by dividing trajectories into 4 blocks and computing the standard error of mean. The folded domains (RRM:2CPE and ZnF:6G99) were maintained as rigid bodies, while the LCD and RGG domains remained flexible. Rigidity was implemented using the fix rigid command in LAMMPS and the hoomd.md.constrain.rigid (Nguyen et al. 2011) function in HOOMD-blue.

### All-atom simulations

Atomistic structures of EWS^LCD^ fragments and EWS^RRM-RGG2^ were generated by a two-step process: first, two representative conformations near the mean radius of gyration (*R*g) for each fragment were selected from CG single-chain simulations which enabled four different combinations of each fragment, then these CG structures were converted to all-atom models using MODELLER (Sali and Blundell 1993). For full-length EWS, we followed the same approach but selected four representative structures. The folded RRM and ZnF were modeled using 2CPE and 6G99 respectively, with the Zn ion modeled as non-bonded to the four coordinating cysteine residues using published parameters (Macchiagodena et al. 2019).

Our system preparation utilized GROMACS v2021.3 (Pall et al. 2020) with the Amber99SBws-STQ force field (featuring improved dihedral corrections for serine, threonine, and glutamine for better representation of helicity in low-complexity regions) (Tang et al. 2020) and TIP4P/2005 (Abascal and Vega 2005) water model. We previously demonstrated that this force field accurately captures the NMR-measured structural features of EWS^LCD^ (Johnson et al. 2024). We placed fragment systems in 13.5 nm octahedral boxes and the full-length protein in a larger 16.5 nm box. After vacuum minimization, systems were solvated with TIP4P/2005 water, followed by energy minimization to relax any steric clashes. Sodium and chloride ions were added to neutralize the charge and reach a salt concentration of 100 mM, using improved salt parameters proposed by Luo and Roux (Luo and Roux 2009). Temperature equilibration was carried out under the NVT ensemble at 300 K using a Nose–Hoover thermostat (Evans and Holian 1985) (coupling constant = 1.0 ps). Pressure equilibration to 1 bar was then performed using the Berendsen barostat (Berendsen et al. 1984) with isotropic coupling (coupling constant = 5.0 ps).

Production simulations were run in Amber22 (Case et al. 2023) at 1 bar and 300 K. The GROMACS topology and coordinate files were converted to AMBER input files (parm7 and rst7) using the “gromber” utility in ParmEd (Shirts et al. 2017). Energy minimization was carried out using the steepest descent and conjugate gradient algorithms, with restraints of 5 kcal/mol/Å^2^ on all heavy atoms. After minimization, we heated the structures by linearly increasing temperature from 0 K to 300 K while applying a force constant of 1 kcal/mol/Å^2^. We then performed an NVT equilibration at 300 K with all restraints removed, followed by an NPT equilibration using a 4 fs time step. The NPT phase employed a Monte Carlo barostat (Åqvist et al. 2004) with a 1.0 ps coupling constant and a Langevin thermostat (Zhang et al. 2017) with a friction coefficient of 1.0 ps^−1^. To enhance computational efficiency, we implemented hydrogen mass repartitioning to 1.5 amu, allowing us to use a 4 fs time step (Feenstra et al. 1999). The SHAKE algorithm (Ryckaert et al. 1977) was used to constrain all bonds involving hydrogen, while long-range electrostatics were treated with the Particle Mesh Ewald (PME) method (Darden et al. 1998) and van der Waals interactions were truncated at 0.9 nm. For each system, we conducted four independent 4 µs simulations, ensuring robust sampling. Based on the radius of gyration autocorrelation analysis, we established a 200 ns equilibration time, which we excluded from subsequent analyses.

To calculate contact maps, we defined a non-bonded contact as any pair of heavy atoms (from residues i and j) within 0.45 nm. The total contact between two residues was computed as the sum of all such heavy atom contacts, capturing side chain contributions, especially from longer residues. Error bars were estimated as the standard error of the mean across the four replicas of each system. For conformational clustering of full-length EWS, we employed the WARIO (González-Delgado et al. 2024) package and visualized results using VMD (Humphrey et al. 1996).

## Author Contributions

EJS: conceptualization (supporting), investigation (lead), formal analysis (lead), visualization (lead), writing – original draft (lead), writing – review and editing (equal). KS conceptualization (supporting), investigation (lead), software (supporting), formal analysis (lead), visualization (lead), writing – original draft (lead), writing – review and editing (equal). LR: investigation (supporting), writing – review and editing (equal). XX: investigation (supporting), writing – review and editing (equal). BF: resources (supporting), supervision (supporting), writing – review and editing (equal). JM: conceptualization (lead), formal analysis (supporting), funding acquisition (lead), resources (lead), software (lead), supervision (lead), writing – review and editing (equal). DSL: conceptualization (lead), formal analysis (supporting), funding acquisition (lead), resources (lead), supervision (lead), writing – review and editing (lead). All authors approve the manuscript for publication.

## Acknowledgements

This study was funded in part by the Marshall County Childhood Cancer Awareness Corporation Alex’s Lemonade Stand Foundation Innovation Award 1335092 and R01GM140127 (DSL), F31CA295030 (EJS), the Welch Foundation A-2113-20220331 (JM), and R35GM153338 (JM). A portion of this work was performed at Brown University with support from the Department of Molecular Biology, Cell Biology & Biochemistry. We gratefully acknowledge the computational resources provided by the Texas A&M High Performance Research Computing (HPRC). This work is based upon research conducted in the Structural Biology Core Facilities, a part of the Institutional Research Cores at the University of Texas Health Science Center at San Antonio supported by the Office of the Vice President for Research and the Mays Cancer Center Drug Discovery and Structural Biology Shared Resource (NIH P30 CA054174).

## Data Availability

NMRPipe processing scripts are available upon reasonable request, expression plasmids were deposited at Addgene: EWS^LCD^ (180467), EWS^1-120^ (180464), EWS^99-199^ (180465), EWS^171-264^ (180466), EWS^RGG1-RRM^ (238369), EWS^RRM-RGG2^ (188046) and EWS^RGG3^ (238370).

## Abbreviations

FUS: Fused in sarcoma
EWS, EWS: RNA-binding protein
TAF15: TATA-binding protein-associated factor 2N
LCD: low-complexity domain
FLI1: friend leukemia integration 1
PARP1: poly [ADP-ribose] polymerase 1
RBD: RNA-binding domain
NMR: nuclear magnetic resonance
RGG: arginine-glycine-glycine
RRM: RNA-recognition motif
ZnF: zinc finger
C_sat_: saturation concentration
CSP: chemical shift perturbations
TEV: tobacco etch virus
MBP: maltose-binding protein
HSQC: heteronuclear single quantum coherence

