## Supporting Figures 1-10 for "Molecular Basis of EWS Interdomain Self-Association and Its Role in Condensate Formation"

† These authors contributed equally

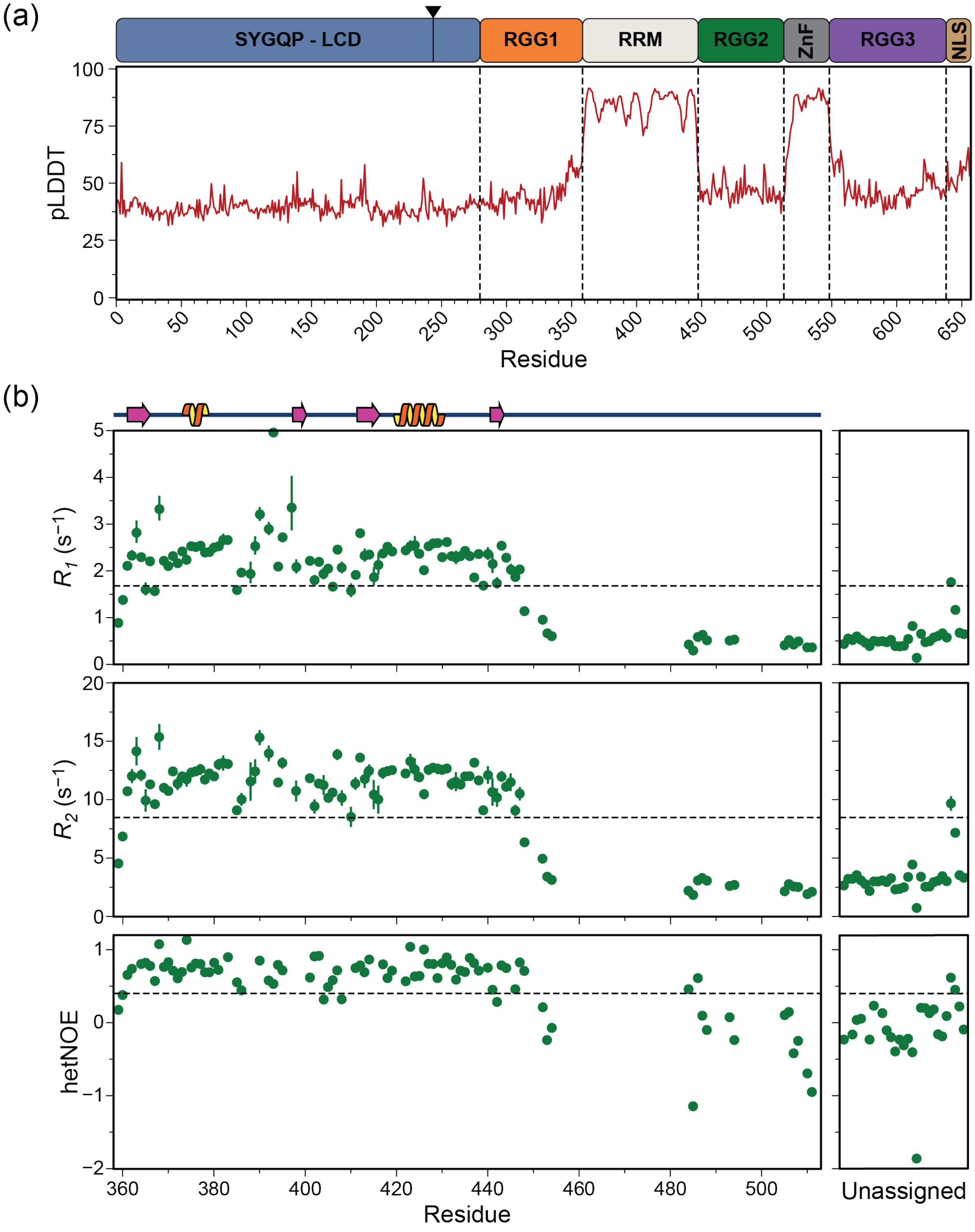


**FIGURE S1.** (a) Predicted local distance difference test (pLDDT) of EWS AlphaFold2 model; pLDDT scores < 50 are considered a very low confidence prediction indicating disorder. (b) The $^{15}$N longitudinal ($R_1$, top), $^{15}$N transverse ($R_2$, middle) relaxation rates, and $^{15}$N heteronuclear NOE values (bottom) measured on 200 µM $^{15}$N EWS$^{RRM\text{-}RGG2}$ in 20 mM potassium phosphate, pH 6, 50 mM KCl, 2 mM TCEP, 1 mM PMSF. 0.4 mM DSS. Secondary structure is depicted above the top panel. Black dashed lines are the average values for $R_1$ (1.67 s$^{-1}$), $R_2$ (8.47 s$^{-1}$) and the hetNOE (0.40).

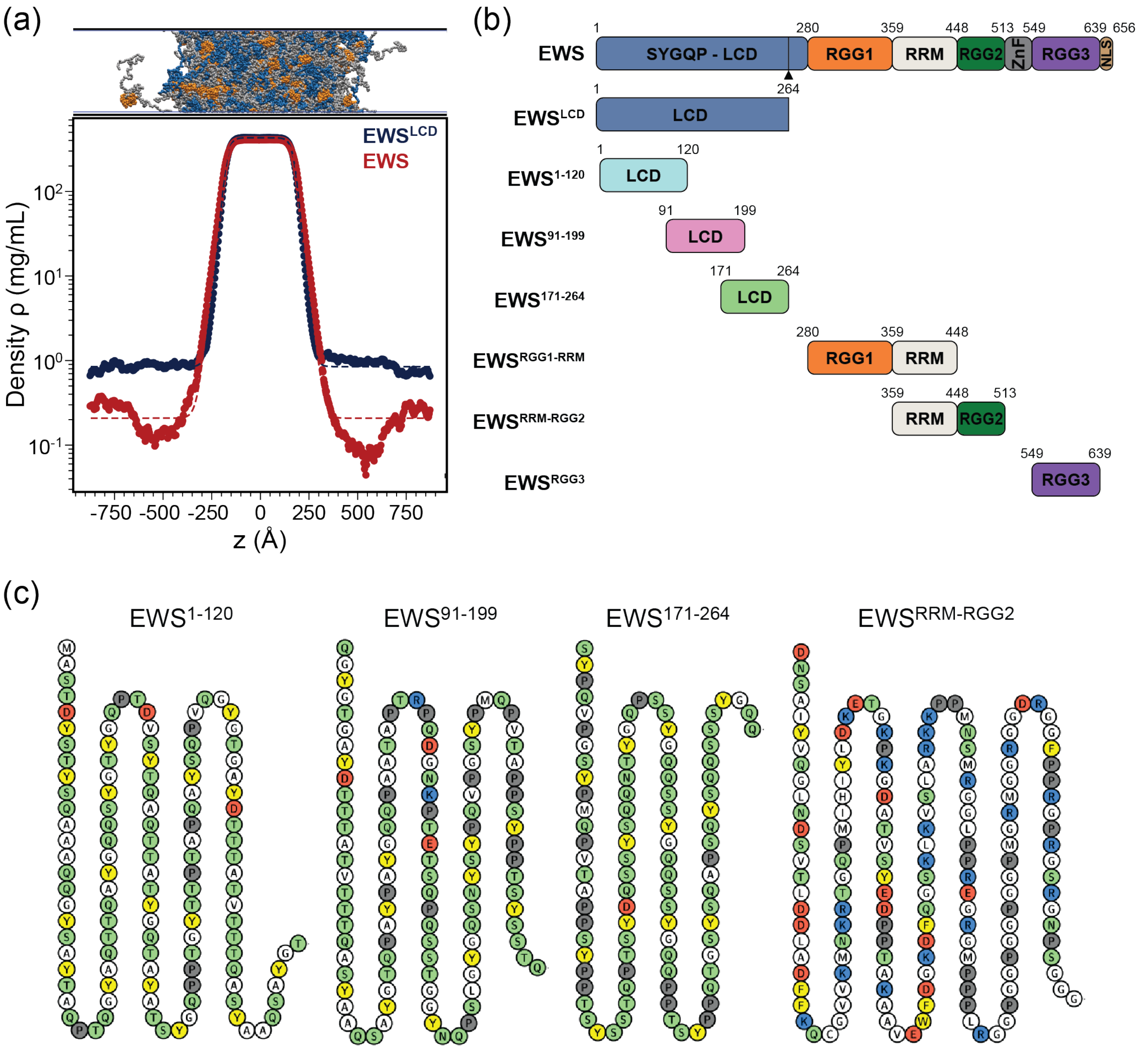


**FIGURE S2.** (a) Density profiles along the z-dimension of the slab geometry for full-length EWS (red) and EWS$^{LCD}$ (blue). (b) Schematic of EWS constructs. Protein sequences and domain margins were extracted from Uniprot and are indicated. Fusion break point is indicated with a triangle. (c) Polypeptide chain sequence for EWS$^{1\text{-}120}$, EWS$^{91\text{-}199}$, EWS$^{171\text{-}264}$, and EWS$^{RRM\text{-}RGG2}$. Residues are color coded as: acidic (red), basic (blue), polar (green), aromatic (yellow), and proline (gray), and others (white).

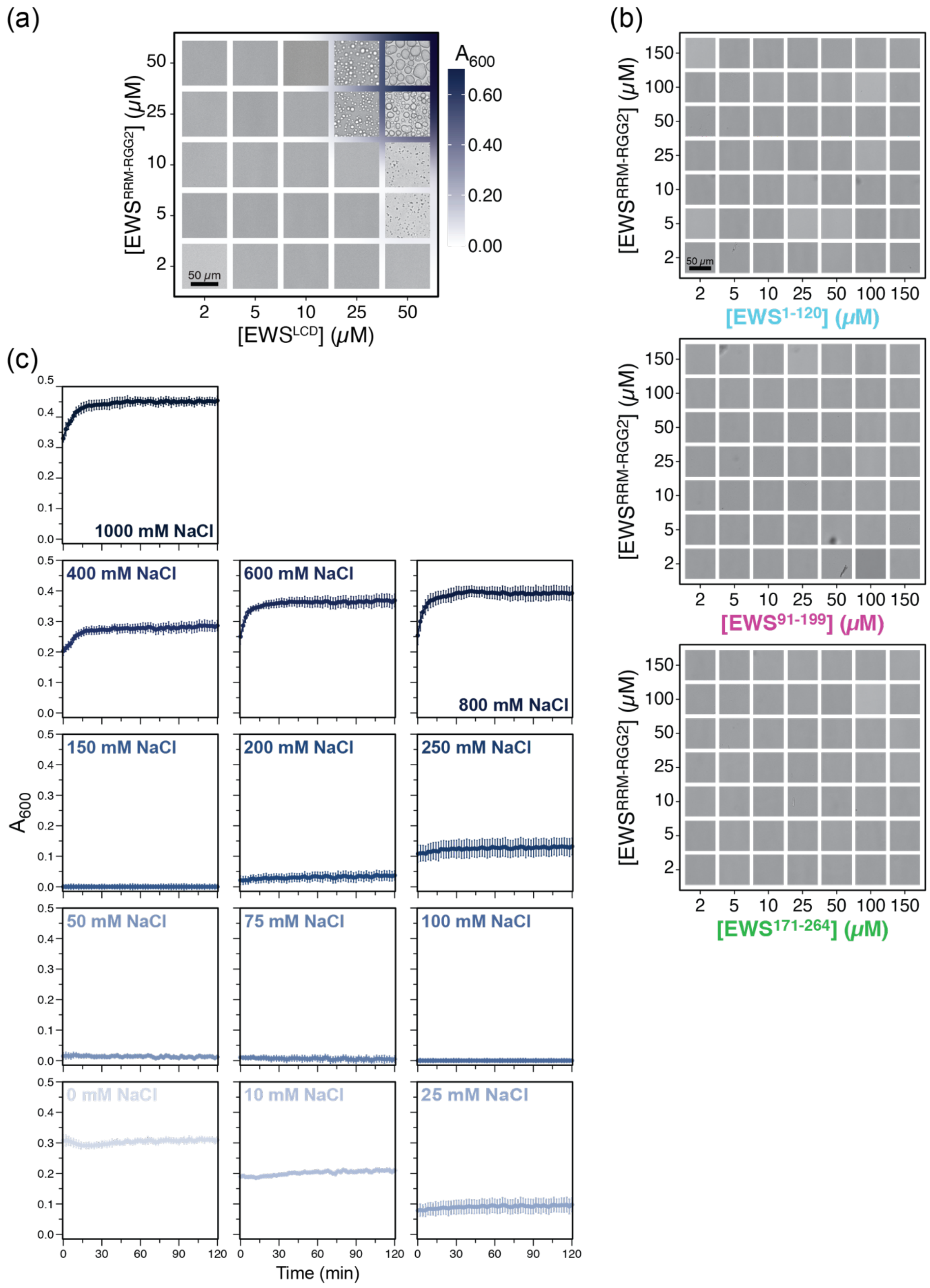


**FIGURE S3.** (a) Phase diagrams of EWS$^{LCD}$ + EWS$^{RRM\text{-}RGG2}$ as a function of protein concentration from bright field images and turbidity measurements. (b) Phase diagrams of EWS$^{RRM\text{-}RGG2}$ + EWS$^{1\text{-}120}$ (top), EWS$^{91\text{-}199}$ (middle), or EWS$^{171\text{-}264}$ (bottom) as a function of protein concentration from bright field images. (c) Turbidity measurements taken over 2 hours of 25 μM EWS$^{LCD}$ + 25 μM EWS$^{RRM\text{-}RGG2}$ across varying NaCl concentration.

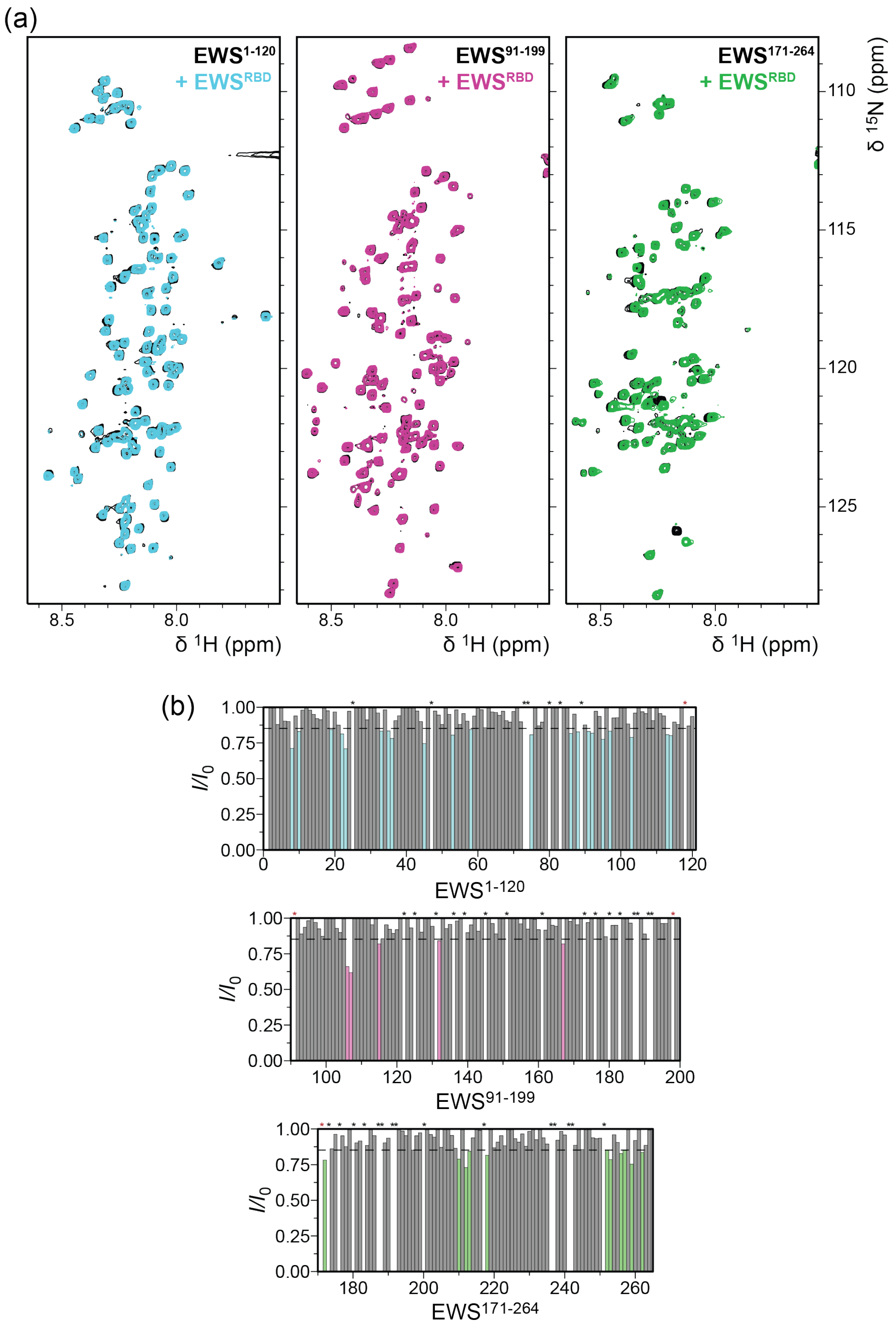


**FIGURE S4.** (a) $^{1}$H,$^{15}$N-HSQC of 100 μM EWS$^{1\text{-}120}$ (left), EWS$^{91\text{-}199}$ (middle), or EWS$^{171\text{-}264}$ (right) (black) and each with a 3:1 molar ratio of EWS$^{RRM\text{-}RGG2}$ (EWS$^{1\text{-}120}$, cyan; EWS$^{91\text{-}199}$, pink; EWS$^{171\text{-}264}$, green). (b) Line broadening of $^{15}$N-EWS$^{1\text{-}120}$ (top), EWS$^{91\text{-}199}$ (middle), or EWS$^{171\text{-}264}$ (bottom) titrated with a 3:1 molar ratio of EWS$^{RRM\text{-}RGG2}$. Residues with intensity differences < 1 SD deviation (dashed black line) are plotted in cyan (EWS$^{1\text{-}120}$), pink (EWS$^{91\text{-}199}$), or green (EWS$^{171\text{-}264}$). Asterisks indicate overlapped/ambiguously (red) or prolines (black).

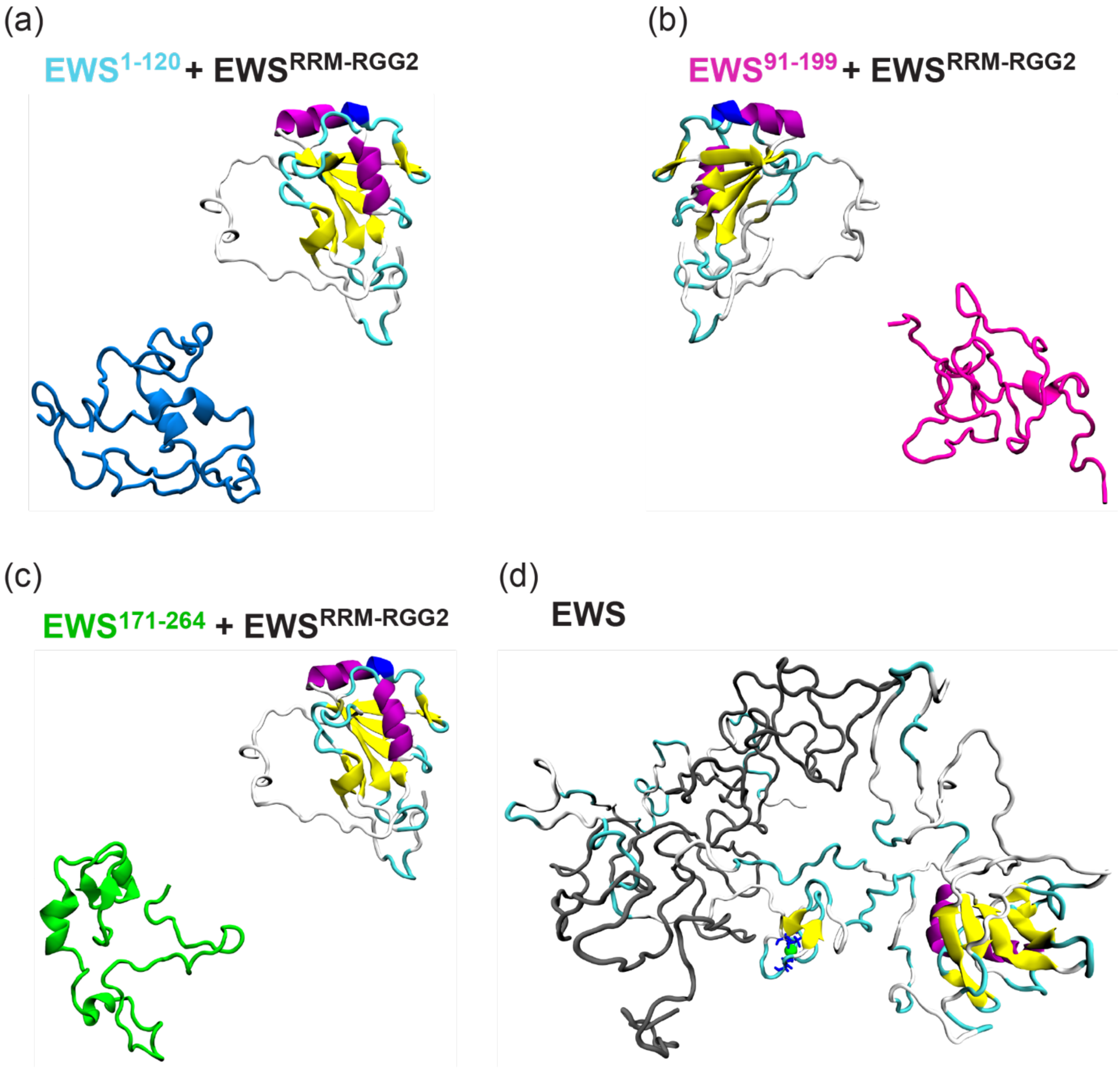


**FIGURE S5.** Representative snapshots from atomistic dilute-phase simulations of $EWS^{RRM\text{-}RGG2}$ paired with (a) $EWS^{1\text{-}120}$ (cyan), (b) $EWS^{91\text{-}199}$ (pink), (c) $EWS^{171\text{-}264}$ (green), and of (d) full-length EWS.

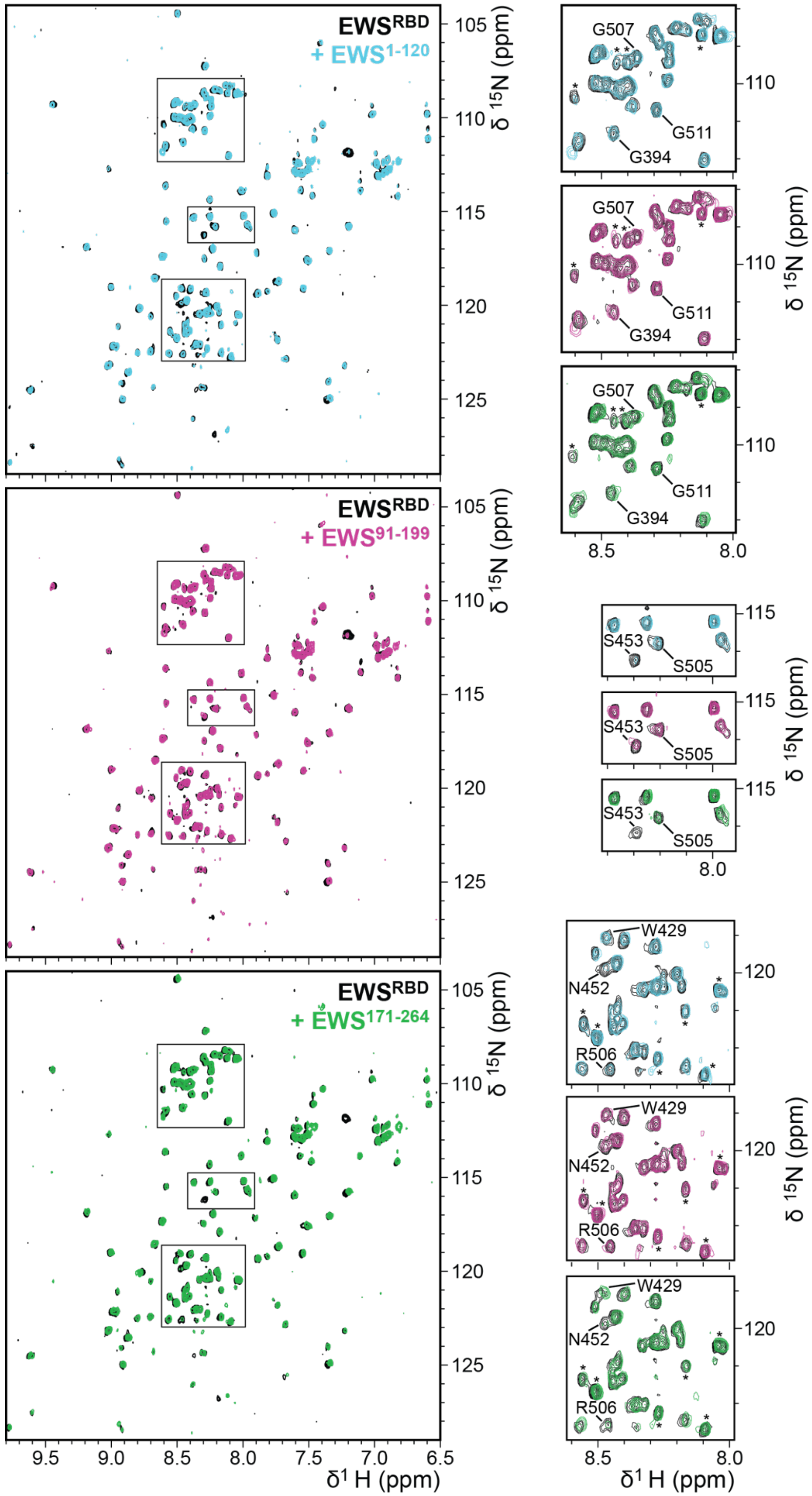


**FIGURE S6.** $^{1}H,^{15}N$-HSQC of 100 μM $^{15}N$-EWS$^{RRM\text{-}RGG2}$ (black) and with a 3:1 molar ratio of EWS$^{1\text{-}120}$ (top, cyan), EWS$^{91\text{-}199}$ (middle, pink), or EWS$^{171\text{-}264}$ (bottom, green). Right panels highlight regions in which peaks are shifted and/or broadened.

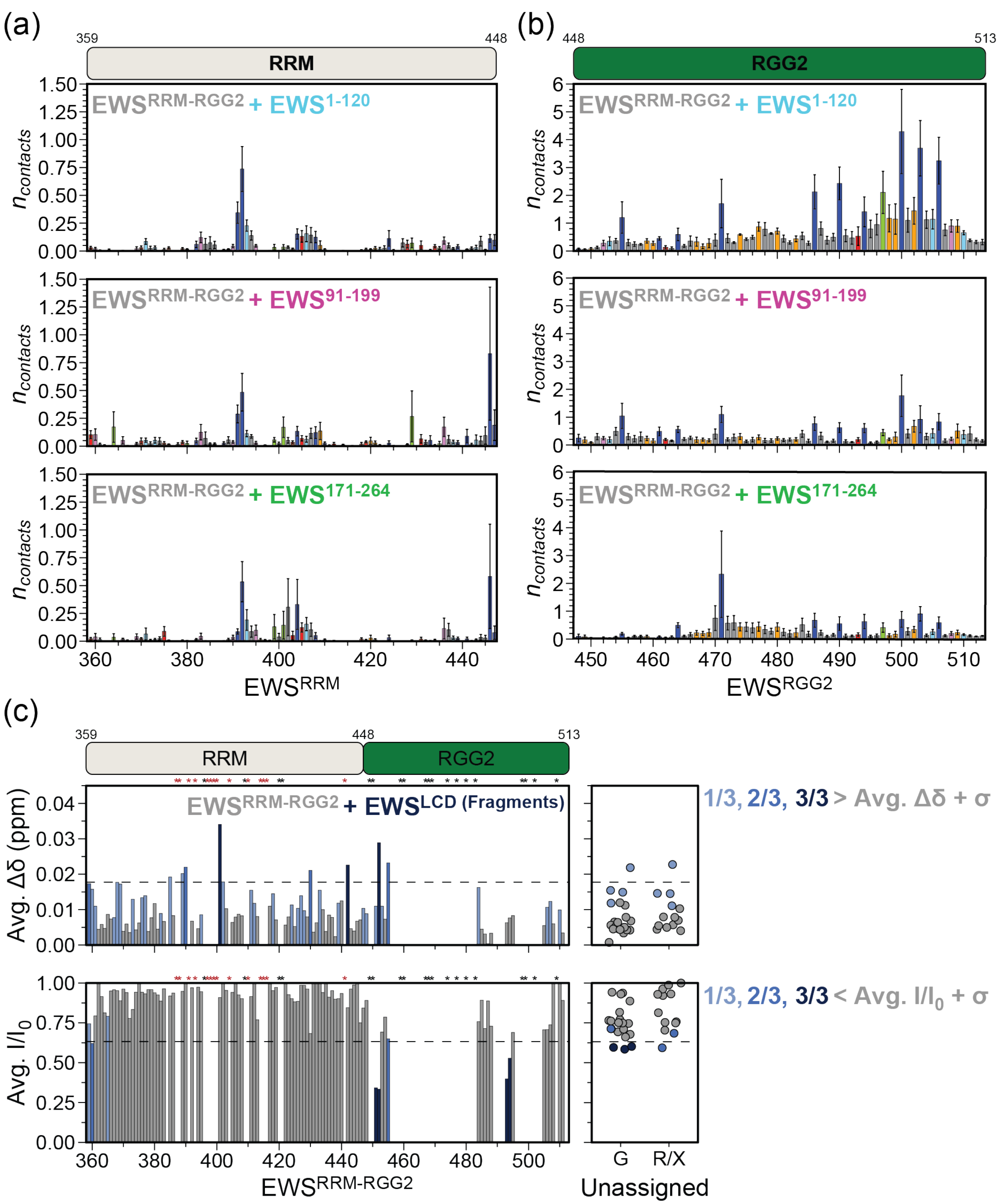


**FIGURE S7.** Average per-residue contacts across (a) EWS$^{RRM}$ or (b) EWS$^{RGG2}$ from atomistic simulation of EWS$^{RRM\text{-}RGG2}$ with EWS$^{1\text{-}120}$ (top), EWS$^{91\text{-}199}$ (middle), or EWS$^{171\text{-}264}$ (bottom). Each residue position is colored by amino acid side chain: acidic (red), basic (dark blue), serine/threonine (light blue), asparagine/glutamine (pink), aromatic (green), proline (orange), and others (gray). (c) Average chemical shift perturbations (CSPs) (top) and line broadening (bottom) of $^{15}$N-EWS$^{RRM\text{-}RGG2}$ titrated with a 3:1 molar ratio of EWS$^{1\text{-}120}$, EWS$^{91\text{-}199}$, or EWS$^{171\text{-}264}$. Residues with CSPs (intensities) < (>) 1 SD (dashed black line) across one, two, or three of the EWS$^{LCD}$ fragments are plotted in light blue, blue, and dark blue, respectively. Asterisks indicate assigned EWS$^{RRM}$ peaks below the limit of detection (red) and from prolines (black). Unassigned glycine (G) or unknown (R/X) resonances are plotted in the right panels.

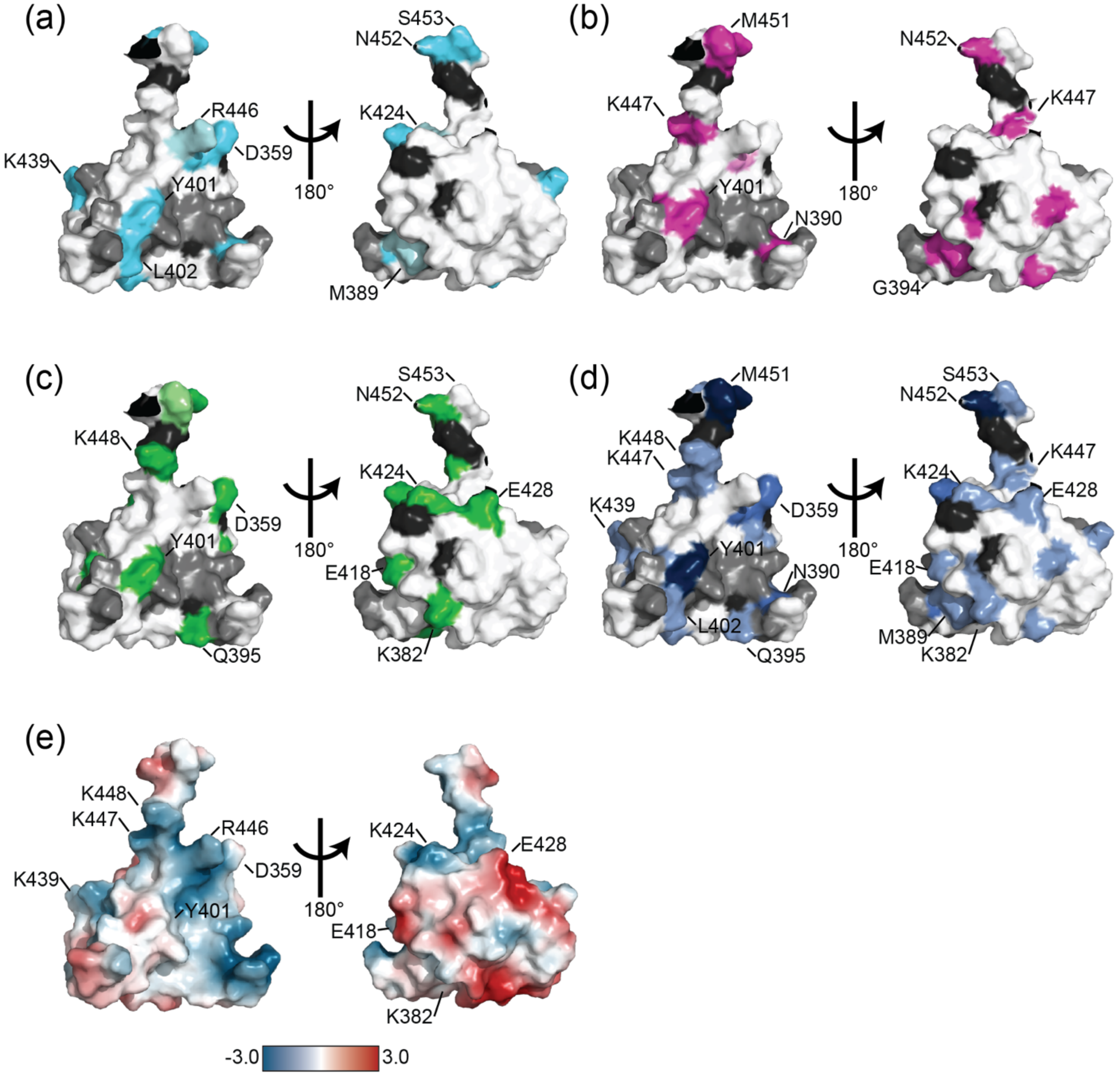


**FIGURE S8.** Residues with CSP (broadening) > (<) than 1SD from $^{15}$N-EWS$^{RRM\text{-}RGG2}$ titrated with a 3:1 molar ratio of (a) EWS$^{1\text{-}120}$ (cyan), (b) EWS$^{91\text{-}199}$ (pink), or (c) EWS$^{171\text{-}264}$ (green) are mapped to the human EWS$^{RRM}$ structure (2CPE). (d) Residues with CSPs (intensities) < (>) 1 SD (dashed black line) across one, two, or three of the EWS$^{LCD}$ fragments are mapped in light blue, blue, and dark blue, respectively. Unassigned/ambiguous and broadened residues are colored dark and light gray, respectively (d) Electrostatic surface of EWS$^{RRM}$ was calculated using the Adaptive Poisson-Boltzmann Solver plugin for PyMol 3.0 (Jurrus et al. 2018).

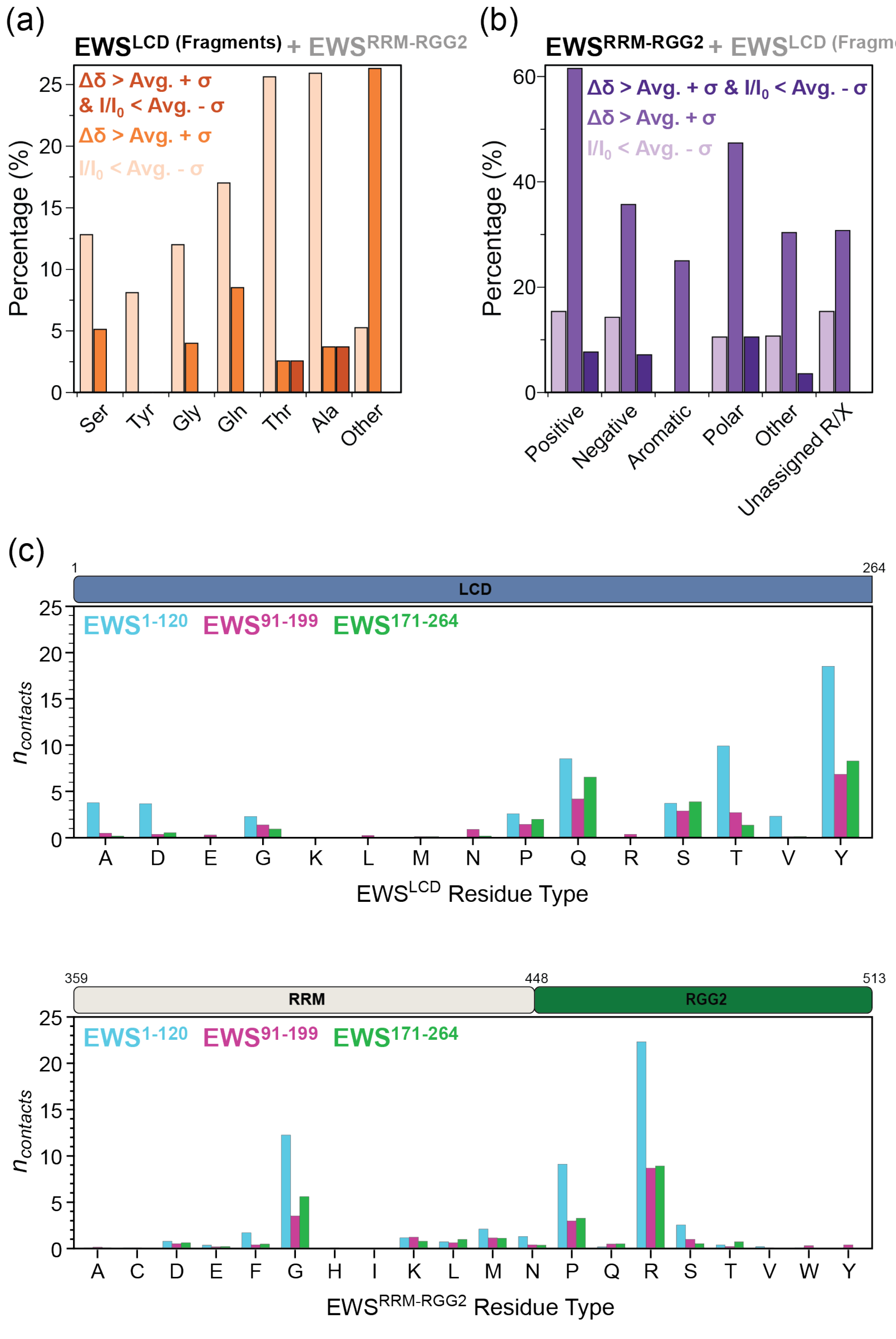


**FIGURE S9.** (a) Percentage of residues with CSPs greater than one standard deviation (orange), intensity differences less than one standard deviation (light orange), or both (dark orange) as a function of specific amino acid for $^{15}$N-EWS$^{LCD}$ fragments titrated with a 3:1 molar ratio of EWS$^{RRM\text{-}RGG2.}$ (b) Percentage of residues with CSPs greater than one standard deviation (purple), intensity differences less than one standard deviation (light purple), or both (dark purple) as a function of residue type for $^{15}$N-EWS$^{RRM\text{-}RGG2}$ titrated with a 3:1 molar ratio of EWS$^{LCD}$ fragments. (c) Average per residue type interdomain contacts formed by EWS$^{LCD}$ (left) or EWS$^{RRM\text{-}RGG2}$ (right) from atomistic simulations of EWS$^{RRM\text{-}RGG2}$ with EWS$^{1\text{-}120}$ (cyan), EWS$^{91\text{-}199}$ (pink), or EWS$^{171\text{-}264}$ (green).

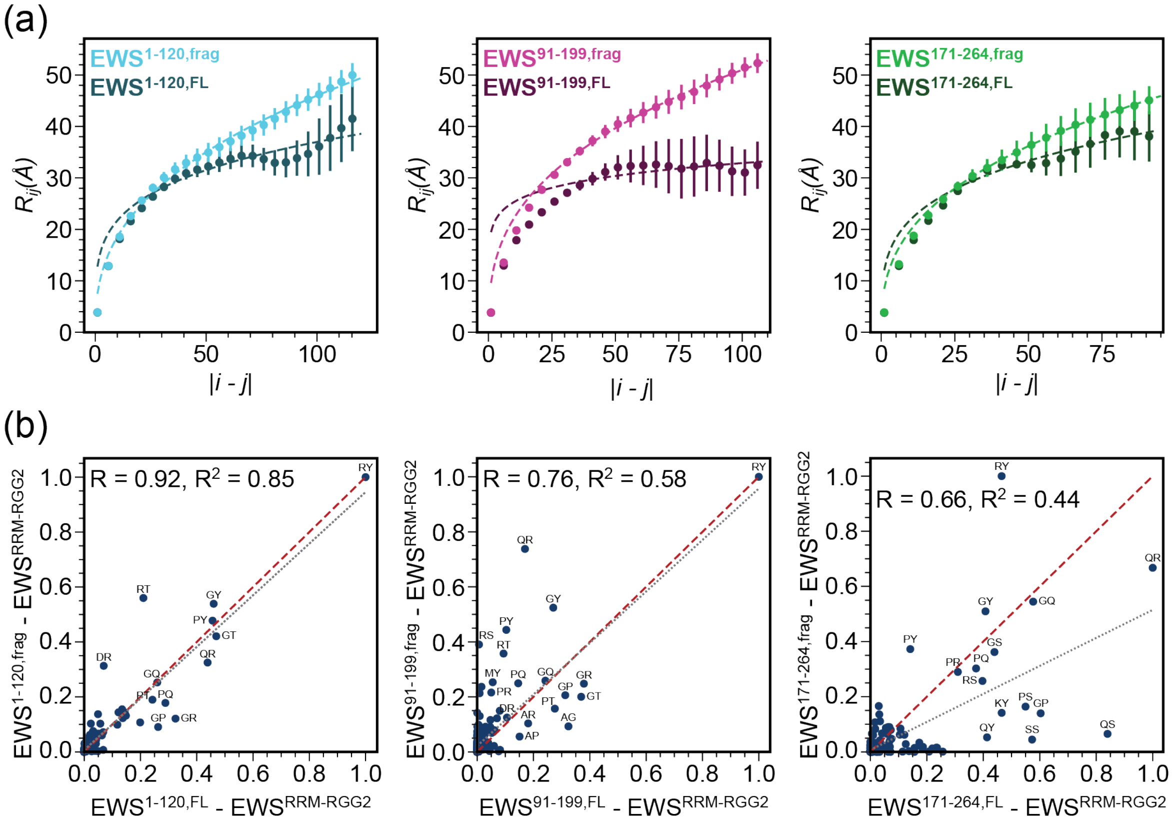


**FIGURE S10.** (a) Inter residue distance plotted as a function of residue separation $|i\text{-}j|$ calculated from atomistic simulations of $EWS^{RRM\text{-}RGG2}$ with $EWS^{1\text{-}120}$ (left, cyan), $EWS^{91\text{-}199}$ (middle, pink), or $EWS^{171\text{-}264}$ (right, green) and single chain atomistic simulations of full-length EWS for each of the same regions (darker shade). (b) Correlation between average residue-type pair contacts in the single chain full-length EWS and $EWS^{LCD}$ fragments for $EWS^{1\text{-}120}$ (left), $EWS^{91\text{-}199}$ (middle), or $EWS^{171\text{-}264}$ (right). The red dashed line represents the diagonal, and the black dotted line indicates the predicted correlation trend.

### SUPPROTING INFORMATION REFERENCES

Jurrus E, Engel D, Star K, Monson K, Brandi J, Felberg LE, et al. Improvements to the APBS biomolecular solvation software suite. Protein Sci. 2018;27(1):112-28. https://doi.org/10.1002/pro.3280
